# The Molecular Interplay of Tau and Motor Proteins on Microtubules

**DOI:** 10.64898/2026.11.02.763819

**Authors:** Yifei He, Julia Peukes, Mert Golcuk, Amanda Jack, Luke S. Ferro, Ruensern Tan, Mert Gur, Eva Nogales, Ahmet Yildiz

## Abstract

Tau is a microtubule (MT)-associated protein essential for neuronal cytoskeletal organization and intracellular transport, yet how tau physically regulates motor proteins remains elusive. Here, we integrate single-molecule imaging, cryo-electron microscopy, and molecular dynamics (MD) simulations to investigate the molecular mechanisms underlying tau-mediated inhibition of kinesin-1 and cytoplasmic dynein motility. We show that tau suppresses MT landing and triggers frequent pausing and premature detachment when motors encounter tau-enriched patches. Unexpectedly, tau impairs each motor using different mechanisms. Tau and kinesin-1 compete for the same surface on tubulin, precluding simultaneous binding to the same MT site. On the other hand, while dynein and tau can co-occupy the same tubulin, tau inhibits dynein’s transition into its high-affinity MT-binding state. Thus, tau can act as a physical or a conformational roadblock in a concentration-dependent manner, exerting distinct structural constraints on kinesin and dynein transport. These findings establish a structural framework for tau-mediated regulation of axonal transport.

## Introduction

Tau is an MT-associated protein (MAP) predominantly expressed in neurons, where it plays a critical role in stabilizing and organizing the MT cytoskeleton and other cellular processes^1^. Tau is particularly enriched in axons and synapses, and its MT binding, organization, and modulation of MT dynamics play important roles in maintaining axonal morphology^2–6^. Its function is tightly regulated by phosphorylation and alternative splicing, which influence tau’s affinity for MTs and its spatial distribution within neurons^1^. Hyperphosphorylation results in dissociation of tau from MTs and formation of neurofibrillary tangles, which is the hallmark of tauopathies, most notably Alzheimer’s disease^7^.

One of the earliest cellular consequences of tau dysregulation is the disruption of axonal transport, leading to synaptic dysfunction and neuronal degeneration^8^. Both overexpression and aggregation of tau cause transport defects, such as impaired delivery of mitochondria, endosomes, and other cargos^9, 10^. Similarly, the knockdown of tau rescues axonal transport in Alzheimer’s models of mouse^11^, indicating that tau plays an inhibitory role in motor-driven transport along axons.

Tau is an intrinsically disordered protein that contains an N-terminal projection region, a proline-rich region, an MT-binding domain (MTBD), and a C-terminal tail (Fig. 1A). The MTBD is composed of four pseudo-repeat sequences (R1–R4), each binding to a tubulin heterodimer. Structural studies showed that tau’s MTBD binds to the outermost ridge on the MT surface in a manner that is compatible with binding up to four adjacent tubulin subunits on the same protofilament^12^. Intrinsically disordered regions in tau are involved in the spacing and bundling of MTs^13^ and liquid-liquid phase separation of tau both in solution and on the MT surface^14–16^.

**Figure 1.**
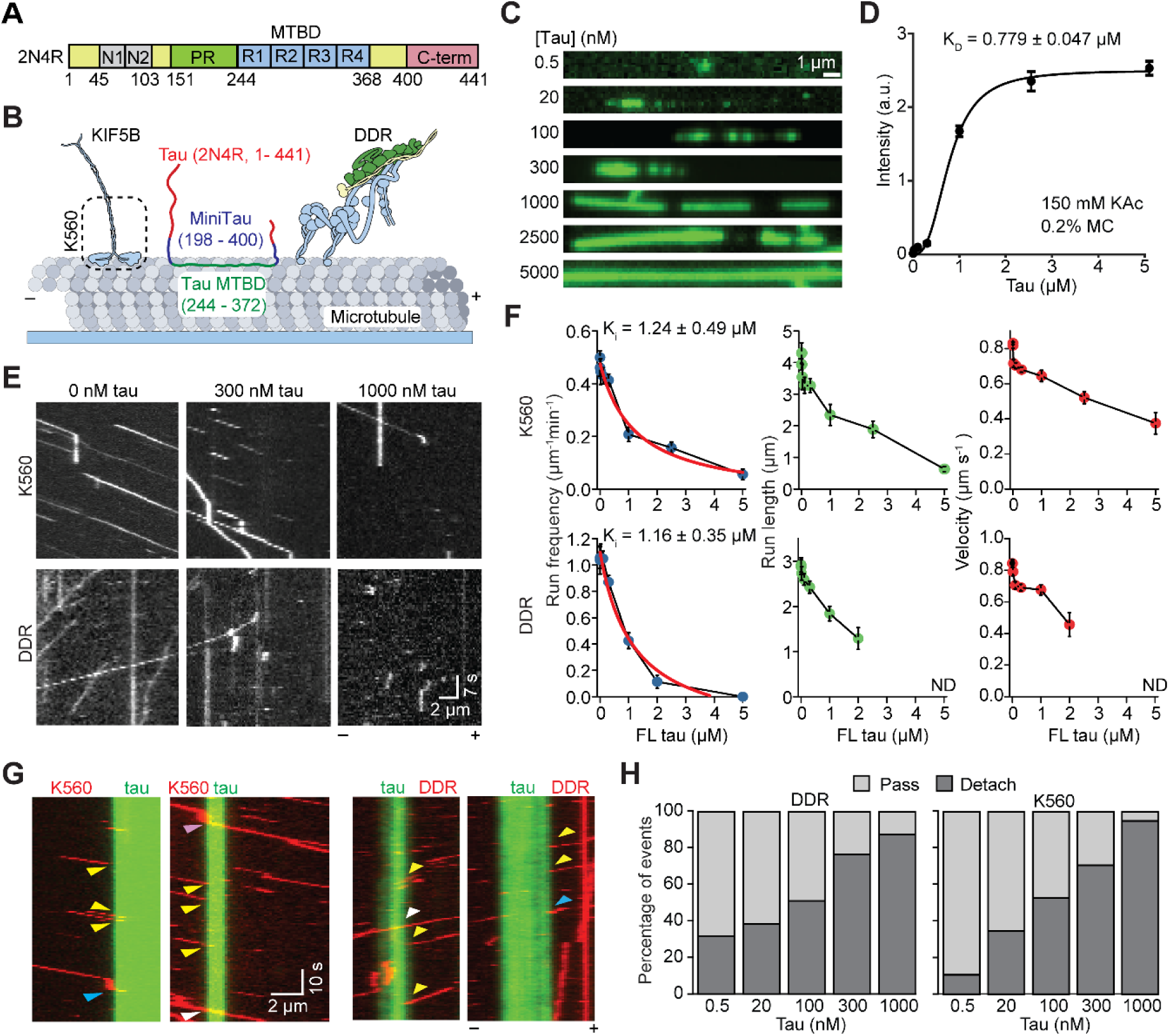
Tau inhibits the motility of both kinesin and dynein. **A.** Domain organization of full-length human 2N4R tau isoform (N1/N2: N-terminal region, PR: proline-rich region, R1-R4: MT binding repeats, C-term: C-terminal region). **B.** Schematic of kinesin and dynein motility on tau-decorated MTs. **C.** Formation of tau envelopes on MTs at different concentrations. **D.** Normalized intensity of FL tau binding to MTs (mean ± S.D.) From left to right, n = 28, 166, 203, 174, 454, 498, 321 MTs. K_D_ (± S.D.) was calculated from a fit to the Hill Equation (solid curve). **E.** Representative kymographs of K560 and DDR on tau-decorated MTs. **F.** Run frequency, run length, and velocity of K560 and DDR at different FL tau concentrations (mean ± s.e.m.). Run frequencies were fitted to an inverse Langmuir isotherm (red curves) to calculate Ki (±SE). From left to right, n = 849, 351, 118, 290, 336, 93, 82, 18 runs for K560 and 502, 453, 455, 492, 551, 206, 209 runs for DDR. **G.** Representative kymographs show K560 and DDR motors pass (white arrowhead), detach (yellow arrowheads), pause and then pass (magenta arrowheads), or pause and then detach (cyan arrowhead) when they encounter tau envelopes. **H.** Percentage of K560 or DDR that pass or detach when they encounter tau envelopes formed at different concentrations. From left to right, n = 19, 75, 95, 242, 60 for K560 and 22, 39, 149, 128, 112 for DDR.

In vitro studies have started to provide structural and mechanistic insight into how tau decoration of MTs regulates motor proteins. Tau reduces the binding affinity and processivity of kinesin-1^15–19^. Unlike kinesin-1, dynein-1 was reported to navigate through tau-decorated regions more effectively^16, 17^, possibly due to its structural differences or distinct stepping mechanism^20^. Differential regulation of these motors raises the possibility that tau acts as a selective filter or modulator of motor traffic, influencing the spatial and temporal delivery of cargo within neurons^21^. However, this model is at odds with the observation that the overexpression of tau halts both retrograde and anterograde transport in axons^8, 22^, and the mechanism behind the regulatory role of tau on motor proteins remains unclear. Tau can physically impede kinesin and dynein motility by competing directly for overlapping binding footprints along the MT^12, 19^. In addition, envelopes formed by tau may function as a selectivity barrier to control access of specific motors to the MT^16,17^. These models are not mutually exclusive, and their relative contributions and distinct mechanistic consequences have not been systematically resolved.

In this study, we performed a comprehensive *in vitro* analysis of how human tau regulates kinesin-1 and dynein-1 motility using single-molecule imaging, cryo-electron microscopy (cryo-EM), and all-atom molecular dynamics (MD) simulations. Motility assays showed that tau inhibits MT landing of both kinesin-1 and mammalian dynein-dynactin-adaptor complexes at high concentrations. At lower concentrations of tau, kinesin and dynein exhibit frequent pausing and MT detachment when they encounter tau envelopes, and kinesin-1 takes more frequent sideways and backward steps when it walks on tau-decorated MTs. Cryo-EM structures of MTs co-decorated with tau and each of these motors revealed that tau and kinesin exclude each other from binding to the same tubulin subunit, whereas tau and dynein can be simultaneously bound to the same tubulin. Instead, MD simulations show that tau inhibits the transition of dynein from a weak to a strong MT affinity conformation through the interaction of a positively charged residue on the dynein MTBD. These results provide new molecular insights into how tau modulates motor-driven transport along MTs.

## Results

### Tau inhibits the motility of both kinesin and dynein

To understand how tau binds MTs, we expressed full-length (FL) human tau (encoded by 2N4R, Fig. 1A) in insect cells and monitored its binding to taxol-stabilized MTs under near-physiological ionic conditions (150 mM Kac). Consistent with previous reports^15, 16^, tau formed liquid envelopes on taxol-stabilized MTs at low (20 nM) concentrations, and these envelopes grew and merged at higher (100 nM) concentrations of tau (Fig. 1B-C). In envelopes, tau intensity per MT length increased at higher tau concentrations (Fig. S1), indicating that MT-bound tau can recruit tau molecules from solution and form a multi-layered tau coat on the MT^23^. To determine the baseline affinity governing initial lattice decoration, we quantified MT binding of tau at different concentrations. Tau binds to taxol-stabilized MTs with a dissociation constant (K_d_) of 779 ± 47 nM (Fig. 1D), substantially higher than the reported K_d_ (60 nM) for insect-cell-derived tau in lower added salt (50 mM)^24^, indicating that electrostatic interactions play a major role in MT binding of tau.

We next tested how tau affects kinesin and dynein motility. Because full-length kinesin is autoinhibited by the folding of its tail onto the motor domain^25, 26^, we expressed a constitutively active human KIF5B construct (K560, amino acids 1-560) that lacks the tail domain (Fig. 1B). We also assembled the dynein-dynactin-BicDR1 (DDR) complex that walks processively along MTs^27^ (Fig. 1B). Previously, *in vitro* studies of MT motors have mostly been performed under low salt conditions, as motors either quickly detach from MTs or exhibit infrequent runs at physiological salt. Because the MT affinity of MAPs is also salt-dependent, MAP-motor interactions have been studied at nanomolar concentrations of MAPs^19^, while cellular MAP concentrations are in the micromolar regime^14^. We addressed this issue by adding 0.1% methylcellulose (a crowding agent), which enabled K560 and DDR motors to take extended processive runs in 150 mM KAc (Fig. 1E-F). These conditions enabled us to investigate the mechanism by which motors are regulated by cellular MAP levels under physiological salt conditions.

MT decoration by tau reduced the run frequency and run length of K560 motors (Fig. 1E-F, Movie S1), as previously reported^8, 15, 17, 28, 29^. Unlike previous reports^16, 17, 29, 30^, tau inhibited DDR motility as strongly as it inhibited kinesin (Fig. 1E-F). The inhibition constants (K_i_) of tau for kinesin (1.2 µM) and dynein motility (1.1 µM) were comparable to each other (Fig. 1F), and slightly higher than the K_d_ of tau for MTs. Because previous studies expressed human tau in *E.coli*^16, 17, 29, 30^, whereas our insect cell-derived tau is phosphorylated at multiple sites and has lower MT affinity^24^, we also compared the two preparations. *E. coli*-derived tau bound MTs more strongly and formed tau envelopes at lower concentrations (Fig. S2A). Similar to insect cell-derived tau, *E. coli*-derived tau also inhibited kinesin and dynein motility at high (>100 nM) concentrations, but K_i_ for kinesin was lower than for DDR, demonstrating that *E. coli*-derived tau inhibits kinesin more strongly than dynein (Fig. S2B-C). These results suggest that the lack of DDR inhibition in previous studies may be due to the low tau concentrations used (∼5 nM)^16, 17, 29, 30^, which are sufficient to inhibit kinesin but not dynein.

Multicolor imaging revealed that both kinesin and dynein had a high probability of passing through tau envelopes formed at low (0.5 - 20 nM) concentrations^15, 16, 30^ (Fig. 1G-H). Although ∼35% of motors detached rapidly upon encountering envelope boundaries, others entered the envelopes and continued moving at reduced velocities before eventually dissociating from MTs (Fig. 1G–H). As tau concentration increased to 300 nM, 60-70% of motors paused or detached when they encountered tau envelopes (Fig. 1D), and this may be due to increased tau density within these regions at higher tau concentrations. Tau envelopes formed at 1,000 nM tau nearly eliminated all motility for both K560 and DDR.

### Tau MTBD is responsible for inhibiting the motors

We next tested whether tau impairs motor motility simply via lattice decoration or through the assembly of cohesive liquid envelopes^15, 16^. If tau envelopes on MTs function as a selectivity filter that excludes motor proteins, MT-bound tau would not be expected to inhibit motor motility under conditions in which tau does not form these envelopes. A minimal tau construct containing only the MTBD (tau-MTBD, amino acids 242-367) does not form liquid envelopes on MTs^16^. Tau-MTBD only weakly interacts with MTs in 150 mM salt and binds to MTs at ∼100-fold lower affinity than FL tau in the absence of added salt (Fig. 2A-C), consistent with a contribution of liquid–liquid phase separation to the stabilization of tau binding to MTs^16^. Importantly, tau-MTBD was sufficient to inhibit both K560 and DDR (Fig. 2D-E and Fig. S3A), demonstrating that neither the N- and C-terminal regions of tau nor the formation of tau envelopes on an MT are necessary for inhibition of kinesin and dynein.

**Figure 2.**
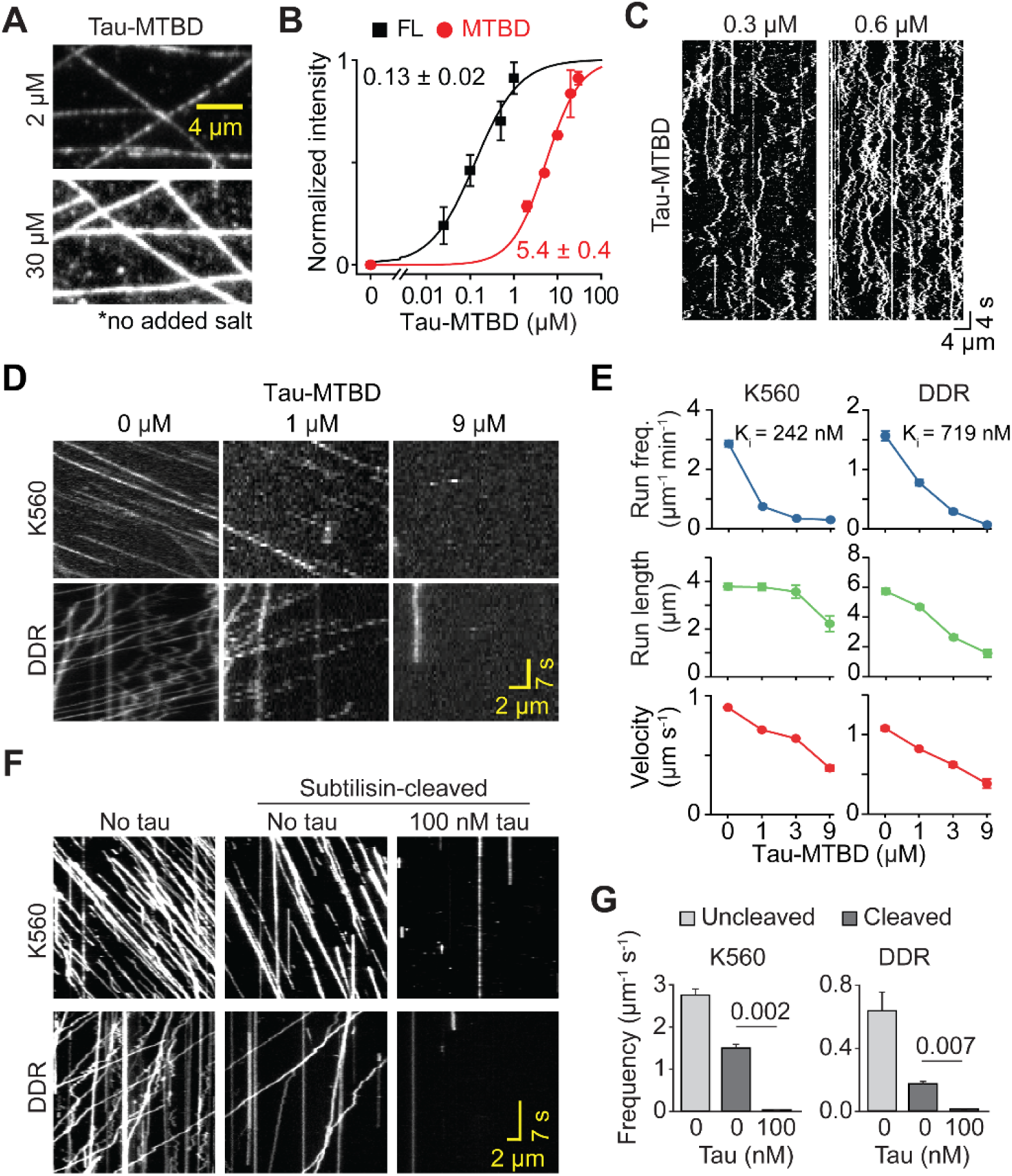
Tau inhibits kinesin and dynein through its MTBD. **A.** Fluorescence images of tau MTBD decoration on MTs. **B.** MT binding affinity of FL tau and tau MTBD in the absence of added salt and methylcellulose (N = 20 per condition, two technical replicates). K_d_ values (±s.e.) were calculated from a fit to the Hill Equation (solid curves). **C.** Kymograph showing the diffusion of fluorescently labeled tau MTBDs on the MT surface. **D.** Representative kymographs of K560 and DDR on MTs decorated with different concentrations of tau MTBD. **E.** Run frequency, run length, and velocity of K560 and DDR at different tau MTBD concentrations (mean ± s.e.m.). From left to right, n = 755, 583, 262, 77 runs for K560 and 445, 596, 167, 24 runs for DDR. Ki values were calculated from a fit to an inverse Langmuir isotherm (see Fig. S3A). **F.** Kymographs of K560 and DDR motility on uncleaved or subtilisin-cleaved MTs in the presence or absence of 100 nM tau. **G.** Run frequency (mean ± s.e.m.) of K560 and DDR motility on uncleaved and subtilisin-cleaved MTs with and without 100 nM tau. From left to right, n = 10, 12, 10 for K560 and 10, 10, 8 for DDR. *P*-values are calculated from a two-tailed t-test.

Previous studies showed that the negatively charged C-terminal tails (CTTs) of tubulin capture motors and retain them on the MT through electrostatic attraction^31, 32^. To investigate whether tau inhibits motors by competing for these CTTs, we cleaved the tubulin tails by subtilisin treatment (Fig. S3B-C). Consistent with previous reports^33, 34^, we observed a 50-70% reduction in the run frequency of kinesin and dynein on subtilisin-treated MTs (Fig. 2F-G). Motors that landed on these MTs walked processively, but the addition of 100 nM tau fully abolished their motility in the absence of CTTs (Fig. 2G-H). We concluded that tau inhibits the binding of kinesin and dynein to the MT surface, rather than preventing their MT recruitment through the CTTs. Collectively, these results suggest that the binding of tau MTBD to the MT lattice is primarily responsible for inhibition of kinesin and dynein motility.

### Tau excludes kinesin but co-exists with dynein on the same tubulin

To determine at the molecular level how tau affects kinesin and dynein binding to MTs, we used cryo-EM to determine the structures of MTs co-decorated with tau and either kinesin or dynein. To minimize potential effects of additional flexible regions on protein recruitment, binding, and affinity that would not be resolved in the structures, we used a previously characterized truncated tau construct containing the pseudorepeats and flanking N-terminal proline-rich region (mini-tau, residues 198-400^16^). Similarly, we used the motor domain of rigor kinesin (K350^E236A^)^35^ and the fusion of dynein MTBD to the seryl-tRNA synthetase (SRS-Dynein_85:82_)^36^, both of which bind MTs with high affinity. Using fluorescence microscopy, we optimized conditions to achieve nearly equal binding of tau and motors under saturating conditions, thereby increasing the likelihood of tau and the motor occupying the same tubulin (Fig. S4).

Based on these concentrations, we prepared cryo-EM grids of peloruside (PelA)-stabilized MTs incubated with mini-tau and either SRS-dynein or rigor kinesin (see Methods; Fig. S5)^37^. Initial average reconstructions of MT segments showed clear density for both tau and dynein or tau and kinesin, indicating that both proteins were associated with the MTs (Fig. 3A). However, because the reconstruction process makes use of the pseudo-helical symmetry of the tubulin dimer on the MT, it averages equivalent positions along the lattice. It thus can artifactually average mutually exclusive MAP and motor densities. To determine true co-binding of a motor and MAP, we used symmetry expansion and signal subtraction (Figs. 3B and S5), followed by focused classification on the region of the bound protein^38^.

**Figure 3.**
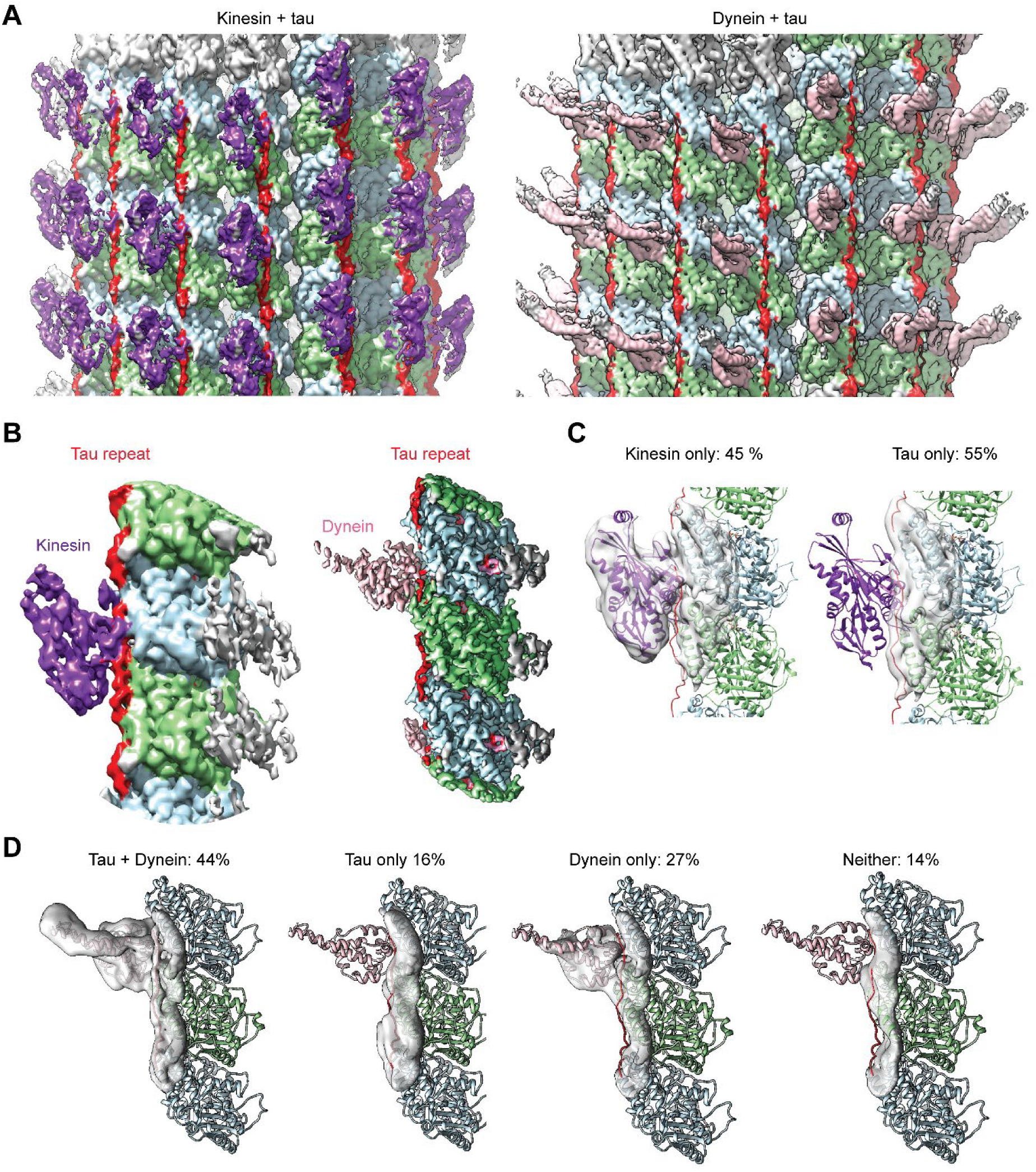
Kinesin and tau exclude each other, whereas tau and dynein can co-occupy the same tubulin dimer. **A.** Cryo-EM reconstructions of MTs decorated with rigor kinesin (purple) and mini-tau (red) or with SRS-dynein (pink) and mini-tau. These reconstructions correspond to the average protein binding density on the MT surface. Surface thresholds were adjusted to visualize substoichiometric tau and motor protein densities. α- and β-tubulin are shown in light green and light blue, respectively. **B.** Reconstruction of an individual tubulin dimer with tau and the respective motor protein, after symmetry expansion and focused refinement. Grey density shows a segment of the adjacent protofilament. **C.** Focused classification after signal subtraction for the kinesin-tau reconstruction showing that one class contains kinesin density only, and the other tau density only, demonstrating that tau and kinesin exclude each other from the same tubulin dimer. **D.** Focused classification for the dynein-tau reconstruction showing four classes with the indicated occupancies. The presence of a class that contains dynein and tau indicates that those two proteins can co-bind the same tubulin dimer. In C and D, fitted structural models were generated from kinesin (PDB: 3J8X)^47^, SRS-dynein (PDB: 6RZB)^45^, R2 tau (PDB: 6CVN)^12^, Pel-A tubulin (PBD: 5SYC^48^; see Methods).

For kinesin and tau, we identified two distinct classes, one with kinesin only and one with tau only. The tau density was in strong agreement with the structure we previously reported for tau on the MT^12^. We did not observe simultaneous binding of tau and kinesin to the same tubulin dimer (Fig. 3C), in agreement with the fact that their footprint on tubulin overlaps. This result shows that tau and kinesin exclude each other due to their overlapping tubulin binding sites^12^ and that tau represents a physical obstacle for kinesin binding to MTs. Unlike kinesin, classification on MTs co-decorated with dynein and tau resulted in four distinct classes: a major class containing tau and dynein co-bound to the MT, two classes of tau or dynein bound individually, and a class without anything bound (Fig. 3D). Our structural data therefore show that, unlike kinesin, dynein and tau can co-bind to the same tubulin dimer, indicating that the tau MTBD does not necessarily constitute a steric roadblock for dynein on the MT surface.

### Kinesin bypasses tau obstacles by backward and sideways stepping on the MT

To understand how kinesin can walk on MTs decorated with tau obstacles, we used single-molecule tracking to analyze kinesin stepping dynamics on tau-decorated MTs at nanometer resolution^39^. We labeled a tail-truncated version of kinesin (K490; amino acids 1-490) with a fluorescent dye at its motor domain and slowed kinesin motility by lowering the ATP concentration to 500 nM, allowing us to resolve individual kinesin steps at 76-ms temporal resolution (Fig. 4A). On undecorated MTs, kinesin mostly follows a single protofilament track, and its motor domains alternately take 16 nm steps in a hand-over-hand fashion^39^. Steps in the sideways or the backwards directions were rarely observed (Fig. 4B-C). At low tau concentrations (<60 nM), kinesin exhibited infrequent sideways or backward steps, with no appreciable change in stepping rate. In contrast, increasing tau concentrations above 100 nM led to a significant increase in both backward- and sideways-stepping events, accompanied by a marked reduction in stepping rate (Fig. 4). These results are consistent with tau acting as a physical obstacle to kinesin motility on the MT. At low tau concentrations, envelopes are rare, and kinesin passes through tau envelopes without impeding its motility. At higher tau concentrations (100–300 nM), kinesins that briefly entered tau envelopes walked by frequently sidestepping to neighboring protofilaments. At higher tau concentrations, the pronounced reduction in kinesin motility in tau envelopes may result from the limited availability of tubulin binding sites on neighboring protofilaments, increasing the likelihood of motor detachment from the MT.

**Figure 4.**
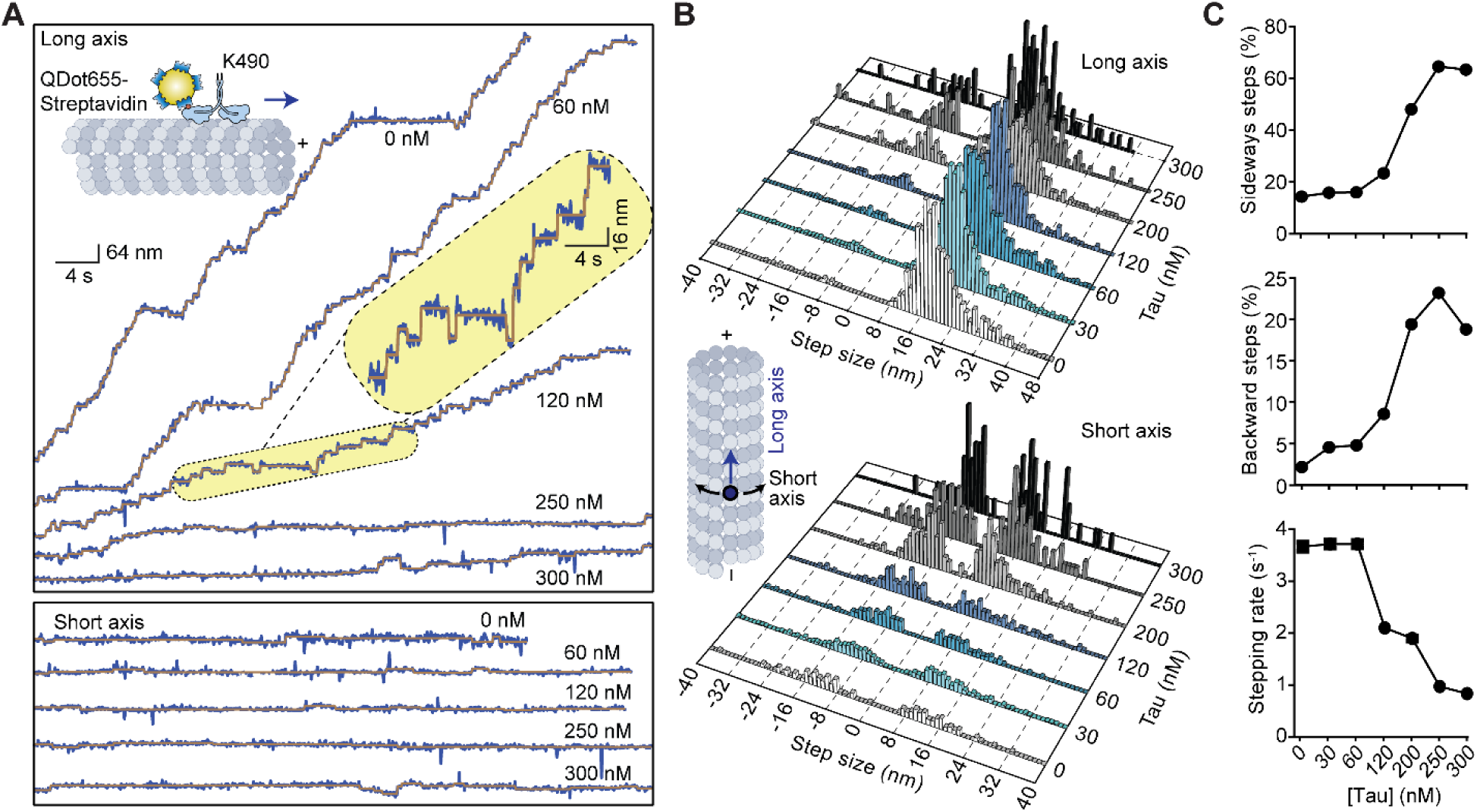
Kinesin frequently takes backward and sideways steps when walking on tau-decorated MTs. **A.** Kinesin K490 was labeled with a bright QDot655 at its motor domain via a biotin-streptavidin linkage. Representative traces of QDot-labeled K490 taking steps along the long (top) and short (bottom) axes of MTs under different tau concentrations. Assays were performed at limiting (500 nM) ATP. Horizontal brown lines represent a fit from a step-finding algorithm. **B.** Kinesin step-size histograms with different FL tau concentrations along the long (top) and short (bottom) axes of MTs. From bottom to top, n = 912, 2120, 911, 744, 479, 198, 101 steps in the long axis and 131, 337, 146, 174, 232, 128, 64 steps in the short axis. **C.** The percentage of steps taken by kinesin in sideways and backward directions, and the stepping rate of kinesin under different concentrations of FL tau on MTs.

We also tested whether multiple motor assemblies walk more successfully on tau envelopes, as this was shown to be an effective strategy to bypass static obstacles on MTs^20, 28, 40^. MT decoration by 50 nM tau at nearly saturating levels fully inhibited the motility of beads driven by multiple K560 or DDR motors as strongly as it inhibited individual motors (Fig. S6). Therefore, teaming up is not an effective strategy for the motors to bypass the high density of tau on the MT, presumably because the motors cannot displace tau stabilized within the envelopes to walk along the MT^15, 16^.

### Tau inhibits MT rebinding of dynein in the weak-affinity state

We next turned our attention to understanding how tau can inhibit dynein motility without directly overlapping with its MT binding site. Our cryo-EM reconstructions using SRS-Dynein_85:82_ captured dynein’s MT-binding domain (MTBD) locked in a high-affinity conformation. However, a processively stepping dynein cyclically alternates between low-affinity (ATP- and ADP·Pi-bound) and high-affinity (apo and ADP-bound) states as it walks^41^. Models of the dynein mechanochemical cycle propose that initial weak engagement of the stepping motor domain with the MT stimulates phosphate release, triggering isomerization into the high-affinity, force-generating state^42^. Tau may therefore selectively disrupt this cycle at the initial landing or transition step, impeding the weak-to-strong affinity switch upon interacting with the MT.

To test this model, we first determined the binding affinities under zero-salt, uncrowded conditions for K350^E236A^ (*K_d_* = 63 *nM*), SRS-Dynein_85:82_ (*K_d_* = 95 *nM*), and mini-tau (*K_d_* = 27 *nM*), identifying conditions yielding near-saturating MT coverage (Fig. S7). We then pre-decorated MTs with 100 nM mini-tau and titrated either K350^E236A^ or SRS-Dynein_85:82_ in the nucleotide-free (apo) state (Figs. 5A and S7A–B). Consistent with our cryo-EM structures, tau and K350^E236A^ segregated into mutually exclusive domains along the MT surface, separated by sharp boundaries^15, 16^. In contrast, tau and SRS-Dynein_85:82_ co-occupied the same MT regions (Fig. 5B). At sub-saturating motor concentrations (5–20 nM), SRS-Dynein_85:82_ engaged tau-decorated MTs far more effectively than K350^E236A^. At saturating regimes (80–400 nM), both motors populated the lattice with efficiencies comparable to undecorated tracks. Crucially, titrating K350^E236A^ progressively displaced mini-tau from the lattice, whereas increasing SRS-Dynein_85:82_ had no measurable impact on mini-tau decoration (Fig. 5C). Reciprocal competition experiments yielded consistent results: when MTs were pre-decorated with 400 nM motor before titrating mini-tau, mini-tau was almost completely excluded from K350^E236A^-coated MTs, but bound efficiently to SRS-Dynein_85:82_-coated tracks. Pre-bound K350^E236A^ and SRS-Dynein_85:82_ remained firmly associated across all mini-tau titrations (Fig. 5D–E), consistent with the exceptionally slow dissociation kinetics characteristic of tight apo-motor binding^43^. Together, these findings demonstrate that high-affinity dynein can stably co-exist with tau on the tubulin lattice, whereas kinesin-1 binding strictly necessitates the physical displacement of tau.

**Figure 5.**
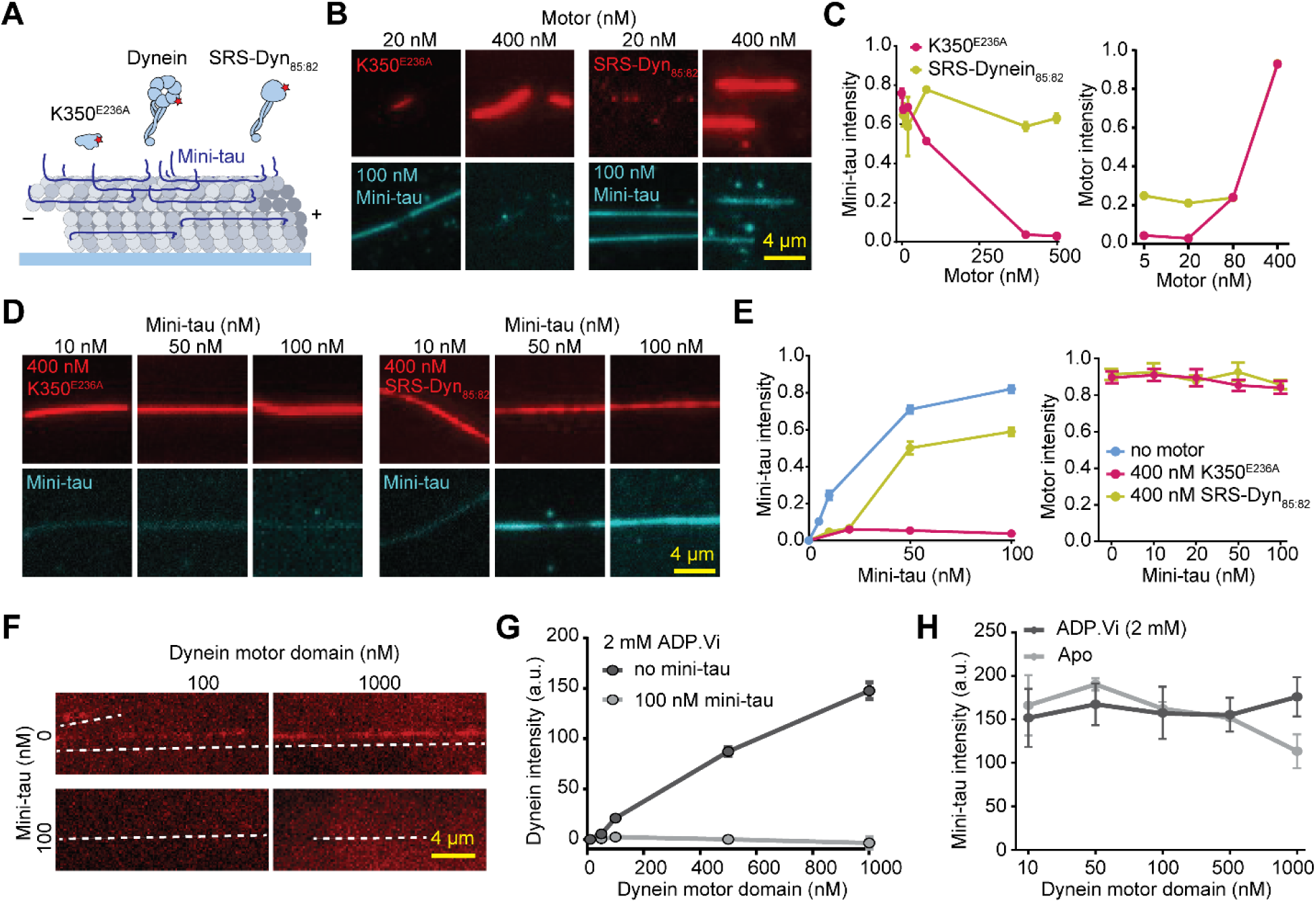
Tau prevents MT binding of dynein in the weak-affinity state. **A.** Schematic showing the binding of monomeric motors to mini-tau-decorated MTs. **B.** MT decoration by 20 nM and 400 nM K350^E236A^ or SRS-Dynein_85:82_ in the presence of 100 nM mini-tau. **C.** (Left) Normalized intensity of 100 nM mini-tau on MTs under increasing concentrations of K350^E236A^ or SRS-Dynein_85:82_. From left to right, n = 65, 47, 162, 54, 63 for SRS-Dynein_85:82_, and 67, 83, 95, 92, 38 for K350^E236A^. (Right) Fluorescence intensities of K350^E236A^ or SRS-Dynein_85:82_ on MTs in the presence of 100 nM mini-tau, normalized to their binding without tau on the MT. From left to right, n = 70, 70, 179, 57 for SRS-Dynein_85:82_, and 63, 77, 95, 104 for K350^E236A^. **D.** MT decoration by 400 nM K350^E236A^ or SRS-Dynein_85:82_ under increasing concentrations of mini-tau. **E.** (Left) Normalized intensity of mini-tau in the presence of 400 nM K350^E236A^ or SRS-Dynein_85:82_. From left to right, n = 77, 66, 91, 92 for no motor, 50, 110, 59, 54 with SRS-Dynein_85:82_, and 156, 54, 94, 92 with K350^E236A^. (Right) Normalized intensities of 400 nM K350^E236A^ or SRS-Dynein_85:82_ at different mini-tau concentrations. From left to right, n = 106, 51, 122, 57, 57 for SRS-Dynein_85:82_, and 102, 98, 58, 97, 104 for K350^E236A^. **F-G.** (Left) Representative images and fluorescence intensity (Right) of the dynein motor domain in the ADP.Vi state in the presence or absence of 100 nM mini-tau. From left to right, n = 53, 78, 51, 65, 68 without mini-tau, and 63, 75, 106, 71, 69 with mini-tau. **H.** Fluorescence intensity of 100 nM mini-tau with increasing concentrations of the dynein motor domain in ADP.Vi and apo states (mean ± S.D.). From left to right, n = 245, 268, 214, 254, 275 in the apo state, and 270, 219, 180, 212, 209 in the ADP.Vi state. In C, E, and G, error bars represent S.E.M.

We also tested if dynein can efficiently interact with tau-decorated MTs in a weak-affinity state. We quantified the MT binding of the motor domain of human dynein-1 on tau-decorated MTs in distinct nucleotide states (Figs. 5F and S7C). To trap dynein in the low-affinity state, we used 2 mM ATP vanadate, which mimics the ADP.Pi state of dynein with weak affinity for MTs^44^. While the dynein motor domain can bind to tau-decorated MTs in the apo states similarly to SRS-Dynein_85:82_, its MT binding was almost fully eliminated by 100 nM mini-tau when in the ADP.Pi state (Fig. 5F-G). Mini-tau intensity remained unaltered across all dynein concentrations regardless of nucleotide state (Fig. 5H), confirming that dynein does not displace tau from the lattice. Together, these results demonstrate that tau does not inhibit dynein motility by steric exclusion, but by selectively precluding MT re-engagement of the stepping head during its transient, low-affinity state.

### Tau prevents docking of the dynein MTBD into the high-affinity state

We performed all-atom MD simulations to gain molecular insight into how tau influences MT binding of dynein when positioned side-by-side on the same tubulin heterodimer. We used the MT-bound structures of the R2 repeat of tau^12^ and dynein MTBD^47^ to build an atomic model of dynein binding to a tubulin heterodimer in the presence of tau (Figure S8A-B). Structural studies have shown that docking of dynein to tubulin is driven by the rotation and translocation of a single helix (H1) towards the cleft between α and β tubulin, while the majority of the MTBD remains relatively rigid upon MT binding^47^. To monitor the docking of H1 to its tubulin binding site in the absence of tau, we used dynein MTBD and a portion of the coiled-coil in the weak-affinity^46^ or the strong-affinity states^47^ and placed them either at the H1 docking site (docked) or 0.7 nm away from this site (undocked; Fig. 6A).

**Figure 6.**
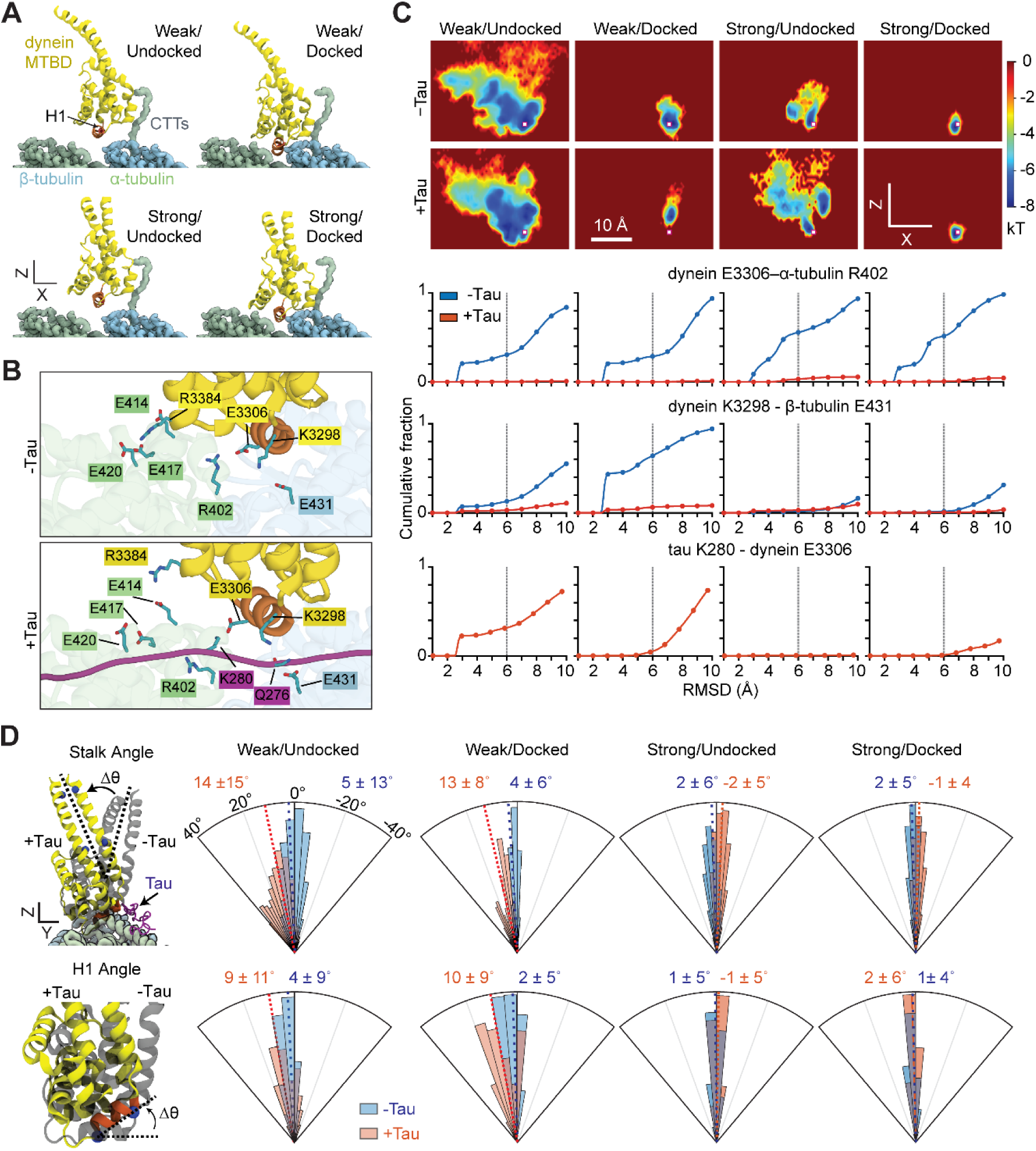
Tau inhibits dynein without overlapping on the MT. **A.** MT docking of the dynein-MTBD was visualized using all-atom MD simulations. Simulations were initiated from weak-(PDB ID: 4RH7)^49^ or strong-affinity (PDB ID: 6RZB)^45^ conformations, either docked or moved 7 Å away from the tubulin binding site (undocked). **B.** Representative snapshots of weak/undocked simulations show that tau disrupts the salt bridge between dynein K3298 and β-tubulin E431 and dynein E3306 and α-tubulin R402. **C.** (Top) Heat maps showing the sampling of H1 near its docking site (white squares) for dynein in weak- or strong-affinity states, starting from docked or undocked positions, and in the presence or absence of tau on the same tubulin. (Bottom) Cumulative fractions of the RMSD distances between key interactions in the presence and absence of tau. **D.** (Top, left) Superimposed snapshots showing that the MTBD of weak-affinity dynein tilts away from tau, whereas the MTBD of high-affinity dynein orients properly as the dynein MTBD attempts to dock onto tubulin. Dashed lines that pass through blue spheres in the stalk define the stalk angle. (Top, right) The angular orientation (Δθ) histogram of the stalk coiled coils with and without tau. The stalk angle in the high-affinity state of dynein (PDB: 6RZB) was set to 0°. (Bottom, left) Superimposed snapshots show that tau causes H1 of weak-affinity dynein to tilt away from its expected orientation as dynein attempts to bind tubulin. Dashed lines that pass through blue spheres in the H1 helix define the angular orientation of H1. (Bottom, right) The angular distribution of H1 (orange) in the presence and absence of tau. H1 orientation in the high-affinity state of dynein (PDB: 6RZB) was set to 0°. Dashed lines represent average orientation (mean ± SD).

In the absence of tau, both weak-affinity and strong-affinity dynein conformers were able to dock onto their tubulin binding site, similar to dynein conformers already docked to tubulin (Fig. 6B-C). The MTBD of dynein formed stable salt bridges with the charged residues on the tubulin surface (Movie S2). In particular, K3298 and E3306 of dynein interacted with E431 of β-tubulin and R402 of α-tubulin, respectively, to lower H1 to the intradimer interface of tubulin (Figs. 6B-C and S8C). In addition, salt bridges between R3384 of dynein and E414, E417, and E420 of α-tubulin appear critical for weak-affinity dynein starting from the undocked conformation to bind near its tubulin binding site. These interactions enabled H1 to position within 0.5 nm root-mean-square deviation (RMSD) from its strong-affinity position in 50% of the total simulation time, and the stalk retained its expected angular orientation relative to the MT (Figs. 6D and S8D).

When we initiated simulations in the presence of the R2 repeat of tau, a weak-affinity dynein from the undocked state failed to lower its H1 into the intradimer interface of tubulin (Figs. 6D and S8C-D). This was because K3298 could not interact with *β*-tubulin E431, and instead, electrostatically trapped via hydrogen bonding with the carbonyl oxygen of tau Q276 (Fig. 6B–C, Movie S2). Similarly, K280 of tau disrupted the salt bridge between E3306 of dynein and R402 of α-tubulin. We also observed a substantial reduction in interactions of R3384 of dynein with E414, E417, and E420 of α-tubulin. This aberrant engagement resulted in deflection of H1’s angular orientation by ∼9° away from the MT surface (Fig. 6D) and tilting of the stalk ∼13° away from tau relative to its native high-affinity orientation (Fig. 6D, Movie S3).

Tau also disrupted salt bridges between dynein E3306 and α-tubulin R402 in weak-affinity dynein already docked to tubulin (Fig. 6B–C) and could not achieve proper stalk and H1 alignment (Fig. 6D). In comparison, even though tau disrupted E3302-R402 interaction in strong-affinity dynein as well, these dynein conformers more frequently formed salt bridges between R3384 of dynein and E414, E417, and E420 of α-tubulin and established additional interactions between their H1 and the tubulin. As a result, strong-affinity dynein docked stably into its binding pocket with H1 and the coiled-coil stalk retaining their canonical conformations (Fig. 6B-D). These results are consistent with our cryo-EM structure showing that dynein MTBD in the strong-affinity conformation can be positioned side by side with tau on the same tubulin dimer (Fig. 3). Thus, our simulations demonstrate that while tau minimally perturbs high-affinity dynein once engaged, it acts as a conformational barrier that precludes weak-affinity dynein from isomerizing into the high-affinity state during initial docking.

## Discussion

Despite the recognized importance of tau in regulating MT-based transport, the molecular mechanisms underlying how tau differentially impacts kinesin and dynein motors have remained controversial. In this study, we integrated single-molecule imaging, cryo-EM, and MD simulations to dissect the mechanisms by which tau modulates the motility and MT engagement of kinesin-1 and cytoplasmic dynein. Our findings reveal that tau is a potent, concentration-dependent inhibitor of both kinesin and dynein motility, consistent with the overexpression of tau to halt both anterograde and retrograde transport in cells. At high concentrations, tau strongly inhibits the landing and processivity of both motors, whereas at lower concentrations, tau induces frequent pausing, detachment, and altered stepping behavior. Importantly, the MTBD of tau alone is sufficient to suppress the motility of both kinesin and dynein. These results suggest that tau-mediated regulation of transport is primarily driven by its occupancy on the MT lattice. Formation of tau envelopes may play additional roles in regulating motor function, as it increases the density and stabilizes the MT binding of tau and potentially reduces the access of the motors to the MT surface.

Although both motors are inhibited by tau, our work uncovers fundamentally different mechanisms by which tau impacts kinesin-1 and dynein. Our cryo-EM structures reveal that tau and kinesin-1 cannot simultaneously bind the same tubulin dimer, establishing a steric exclusion mechanism. This stands in contrast to a previous cryo-EM study proposing co-occupancy between kinesin-1 and the related MAP4^46^. Here, focused 3D classification resolved this discrepancy: standard helical reconstruction averages tubulin dimers across the lattice, creating the false appearance of co-binding from an ensemble of mutually exclusive binding events. By circumventing this averaging artifact, our classification analysis confirms that tau and kinesin-1 never occupy the same tubulin subunit concurrently.

Unlike kinesin, dynein and tau can simultaneously bind to the same tubulin dimer on the MT, indicating that tau’s inhibitory effect on dynein is not due to direct competition for binding sites. Instead, our MD simulations suggest that tau interferes with a key salt bridge between the dynein MTBD and β-tubulin that facilitates the transition of dynein into its high-affinity MT-bound state. In the presence of tau, this salt bridge cannot form, and dynein is prevented from properly docking onto the MT surface. This form of inhibition represents a distinct mechanism of motor regulation: allosteric reduction of binding affinity through the modulation of specific electrostatic interactions at the motor–MT interface.

Our single-molecule stepping assays further highlight the adaptability of kinesin in the face of tau obstacles. On tau-decorated MTs, kinesin exhibits an increased frequency of sideways and backward stepping, consistent with a bypass mechanism in which the motor navigates around tau by switching protofilaments. Tau also decreases the overall stepping rate of kinesin, which may be related to the rebinding of kinesin to the same tubulin if the next tubulin site on the same protofilament is occupied by tau. However, at high tau densities, kinesin’s ability to bypass is severely compromised, resulting in stalling or MT detachment. Interestingly, both the ability of dynein to take frequent sideways steps on the MT^20^ or the cooperative behavior of multiple kinesins or dyneins fail to rescue motility under high tau conditions, indicating that the lack of accessible tubulin binding sites on neighboring protofilaments limits the motor’s options, leading to transport arrest.

Previous in vitro studies proposed that tau selectively inhibits kinesin but not dynein^16, 17, 29, 30^. These studies often used a bacterially expressed tau construct that lacks phosphorylation and binds MTs with nanomolar affinity under low salt conditions. In comparison, we used tau expressed in insect cells, which carries 5-10 phosphorylation modifications and has a weaker affinity to bind MTs in physiological salt^24^. We showed that *E. coli*-derived tau inhibits kinesin more strongly than dynein, but similar to insect cell-derived tau, it inhibits both kinesin and dynein at high concentrations. It remains to be determined whether phosphorylation plays a specific role in regulation of MT motors.

Our MD simulations indicate that tau impedes docking of the dynein MTBD in the weak-affinity conformation, providing an explanation for the reduced MT binding observed in the ADP.Pi state. However, in the presence of tau, dynein can still bind to the MT when it switches to a high-affinity state after releasing the inorganic phosphate (ADP-bound state). These results provide an explanation for how tau inhibits dynein motility. During processive motility, the stepping head releases from the MT upon ATP binding and remains unbound in the ADP.Pi state. Solution kinetic studies proposed that initial weak engagement of the stepping head with the MT accelerates phosphate release^42^, promoting its MT binding. Tau may disrupt this interaction, increasing the time the stepping head spends in the one-head-bound state. In this state, the motor becomes vulnerable to MT detachment due to the nucleotide turnover of the MT-bound head and prematurely ends its processive run.

We propose that tau exhibits concentration-dependent, mechanistically distinct inhibition of both anterograde and retrograde transport along axons, rather than acting as a directional filter that preferentially permits the retrograde motor dynein. Tau exerts inhibitory effects via distinct mechanisms: steric exclusion for kinesin and competition for MT-binding interactions for dynein. These findings unify disparate models of tau regulation and offer a new mechanistic explanation for how tau dysfunction impairs intracellular transport in neurodegenerative disease. Future studies are required to explore how post-translational modifications and disease-associated mutations of tau affect its regulatory role in motor-driven transport.

## Acknowledgments

We thank R. McKenney for tau constructs, P. Northcote (Victoria Univ. of Wellington) for providing Peloruside A, J. Slivka, Y. Zhao and J. Fernandes for assistance with protein purification, J. Zhu and J. Flanagan for assistance with fluorescence image acquisition and processing, N. Kambdur and H. Adams for assistance with cryoEM sample preparation and optimization, Q. Fang for help with initial cryoEM data analysis, P. Grob for electron microscopy and computational support, the Cal-Cryo facility at UC Berkeley for EM imaging, and Indiana Jetstream2 and NCSA Delta for MD simulations. This work was supported by grants from the National Institute of General Medical Sciences (GM136414 (A.Y.), GM127018 (E.N.), GM123655 (L.S.F.), and the National Science Foundation (MCB-2534447, A.Y.; DGE-1752814, A.J.), Advanced Cyberinfrastructure Coordination Ecosystem (ACCESS, BIO240144, M.Gur). E.N. is a Howard Hughes Medical Institute Investigator.

## Author contributions

A.Y., E.N., and M. Gur conceived the project. Y.H., A.J., L.S.F., and R.T. and purified the proteins, performed single-molecule experiments, and analyzed the data. J.P. performed cryo-EM sample preparation, data collection, and analysis. M. Golcuk and M.Gur performed model building, MD simulations, and analysis. Y.H., J.P., M. Golcuk, M. Gur, E.N., and A.Y. wrote the manuscript.

## Supplementary Tables

**Table S1.** The list of constructs used in this study.

| Name | Description | Source | Expression | Figures |
|---|---|---|---|---|
| FL tau | Human ybbR::tau<sup>1-441</sup> | This study | Sf9 | 1, 2, 4, S1, S6 |
| Mini-tau | Human StrepII::sfGFP::tau<sup>198-400</sup> | Tan et al. (2019)<sup>16</sup> | E.coli | 3, 5, S4, S5, S7 |
| Tau MTBD | Human StrepII::sfGFP::tau<sup>244-372</sup>::FLAG tag::His | Tan et al. (2019)<sup>16</sup> | E.coli | 2 |
| FL tau | 2N4R Tau<br>6xHis:: TwinStrepII:: sfGFP:: 2N4R | Tan et al. (2019)<sup>17</sup> | E.coli | S2 |
| K560 | Human KIF5B<sup>1-560</sup>::GFP:: SNAP | Ferro et al. (2022)<sup>38</sup> | Sf9 | 1, 2, S6 |
| K490 | Human KIF5B<sup>1-490</sup>::GFP:: SNAP | Ferro et al. (2022)<sup>38</sup> | Sf9 | S2 |
| K490 E215C | Human KIF5B<sup>1-490</sup> E215C (Cys-light) | This study | Sf9 | 4 |
| K350<sup>E236A</sup> | Human KIF5B<sup>1-350</sup>::His tag (E236A) | This study | E.coli | 3, 5, S4, S5, S7 |
| Phi mutant dynein | SNAPf::DYNH1C1<sup>K1610/R1567E</sup>-IC2C-LIC2-Robl1-Tctex1-LC8 | Schlager et al. 2014<sup>50</sup> | Sf9 | 1, 2, S2, S6 |
| Human dynein-1 motor domain | SNAPf::DYNH1C1<sup>(Δ1-1289)</sup> | This study | Sf9 | 5, S7 |
| SRS-Dynein<sub>85:82</sub> | The distal portion of the stalk of mouse dynein-1 fused to SRS from *T. thermophilus* | Carter et al. (2008)<sup>51</sup> | E.coli | 3, 5, S4, S5, S8 |
| BicDR1 | Mouse His::ZZ::TEV::BicDR1<sup>1-577</sup>::SNAP | Ferro et al. (2022)<sup>38</sup> | Sf9 | 1, 2, S2 |
| BicD2N | Mouse His::ZZ::TEV::BICDN<sup>1-400</sup>::GFP | Schlager et al. (2014)<sup>51</sup> | Sf9 | S6 |

**Table S2.** Cryo-EM dataset and model-building statistics.

| **Sample** | **MTs + tau/kinesin** | **MTs + tau/dynein** |
|---|---|---|
| Microscope | Talos Arctica | Talos Arctica |
| Voltage (kV) | 200 | 200 |
| Camera | K3 | K3 |
| Defocus range | −1 to −2 µm | −1 to −2 µm |
| Data collection software | SerialEM | SerialEM |
| Frames | 50 | 50 |
| Total dose (e⁻/Å²) | 50 | 50 |
| Pixel size (Å/px) | 1.14 | 1.14 |
| Super-resolution mode | Yes | Yes |
| No. of movies | 2,315 | 3,200 |
| No. of CTF-filtered movies | 2,190 | 2,910 |
| No. of initial particles | 108,044 | 112,292 |
| No. of 14-PF particles | 75,740 | 65,769 |
| Resolution after seam finding (Å) | 3.53 | 3.49 |

**Table S3.**
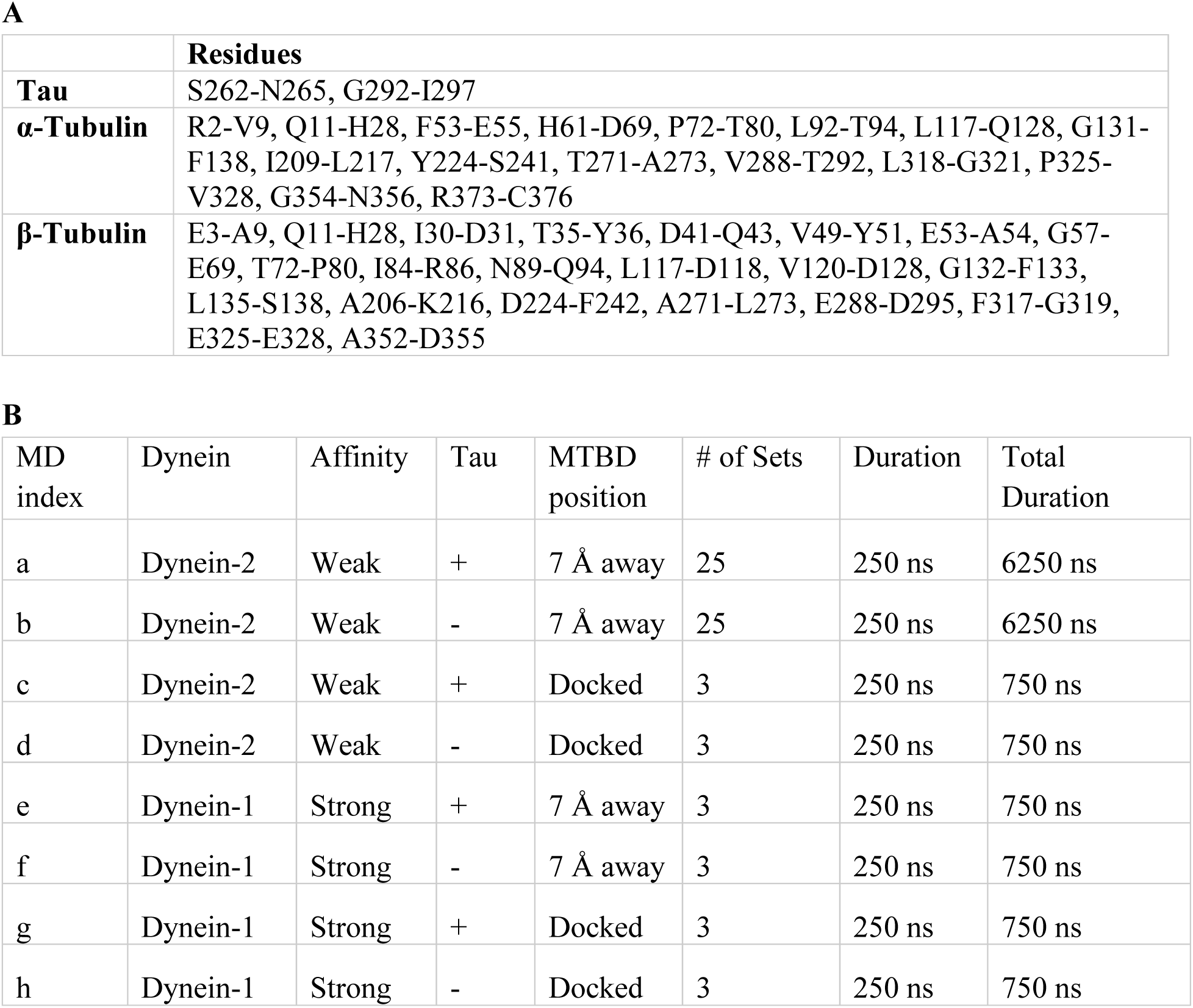
Constraints and conformations used in MD simulations. **A.**The list of amino acids from terminal residues of the R2 segment of tau and residues of α- and β-tubulin that face the MT lumen with constrained C_α_ atoms during MD simulations. Constraints were applied to residues at both termini of the tau R2 segment to ensure tau remains bound to tubulin. Additional constraints were applied to residues facing the MT lumen in tubulins to mimic the protofilament environment and preserve the structural integrity of the MT lattice. **B.** The list of MD simulations performed in this study. Dynein-2 and dynein-1 MTBD conformers represent weak and strong MT-binding affinity, respectively. These conformations were simulated in the presence (+) or absence (-) of the tau R2 segment on tubulin. For each condition, the MTBD was positioned either in its docked position on the tubulin or 7 Å away from the binding site. Each independent simulation was performed for 250 ns, a total of 17 μs.

## Supplementary Figures

**Fig. S1.**
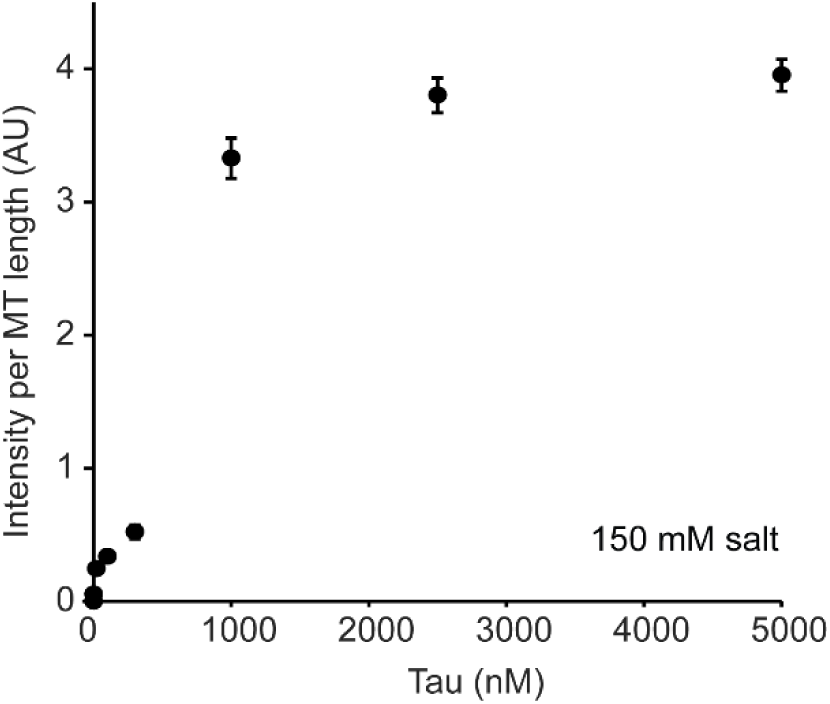
Tau intensity grows higher in envelopes formed at higher concentrations. Fluorescence intensity per MT length (mean ± SD) for tau envelopes formed at different tau concentrations. From left to right, n = 19, 61, 51, 62, 62, 59, 48 envelopes.

**Fig. S2.**
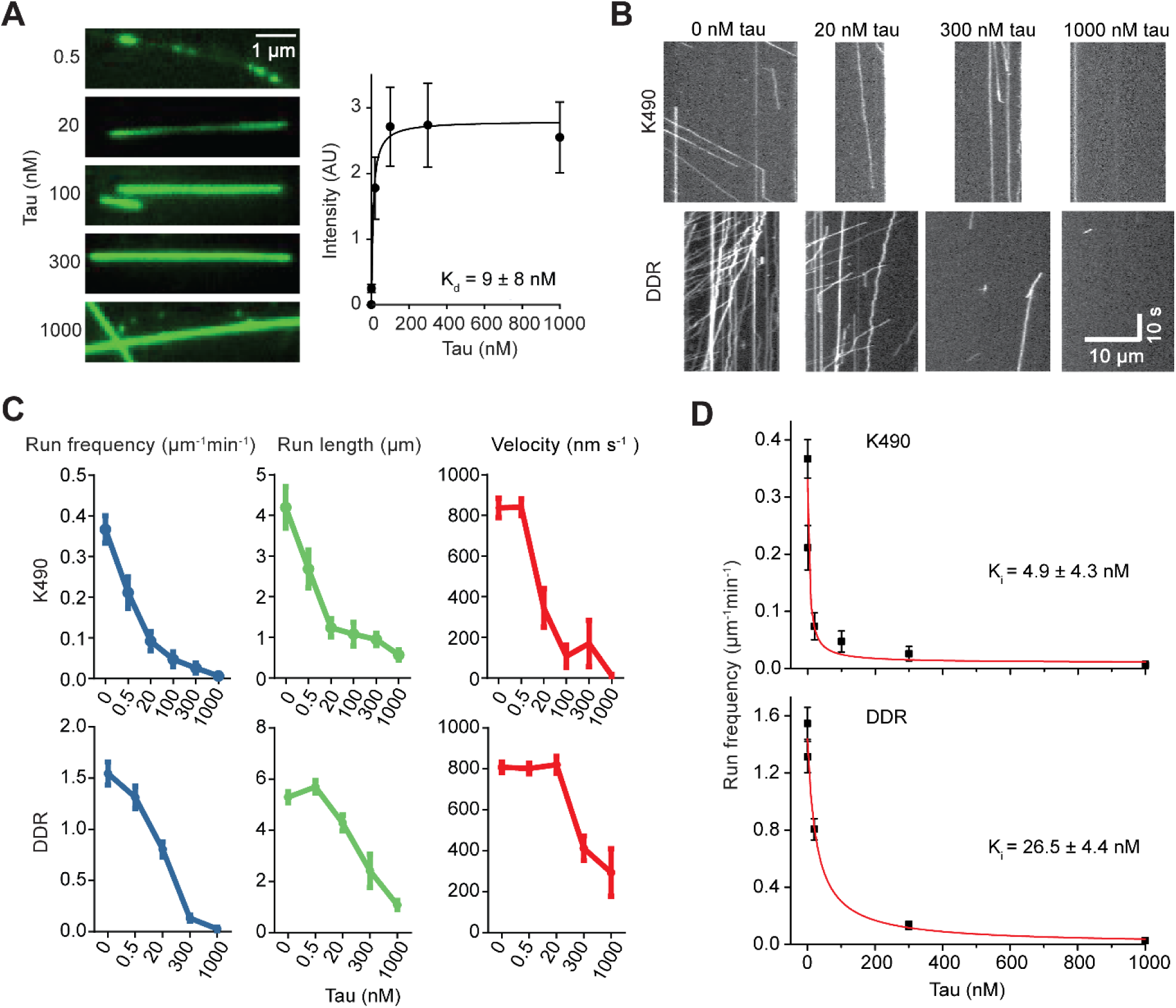
MT binding and motor regulation of tau expressed in E. coli. **A.** (Left) Formation of tau envelopes on MTs at different concentrations. (Right) Fluorescence intensity of FL tau binding to MTs. From left to right, N = 31, 40, 48, 49, 27 MTs. The dissociation constant (± S.D.) was calculated from a fit to the Hill Equation (solid curve). **B.** Kymographs of K490 and DDR with and without tau purified from E. coli. **C)** The run frequency, run length, and velocity of K490 and DDR with and without tau purified from E. coli (mean ± s.e.m.). **D.** Run frequency (mean ± s.e.m.) of K490 and DDR was fitted to an inverse Langmuir isotherm (red curves) to calculate K_i_ (±SE). In C and D, from left to right, n = 49, 42, 15, 11, 8, 2 runs for K490 and 295, 345, 155, 34, 6 runs for DDR.

**Fig. S3.**
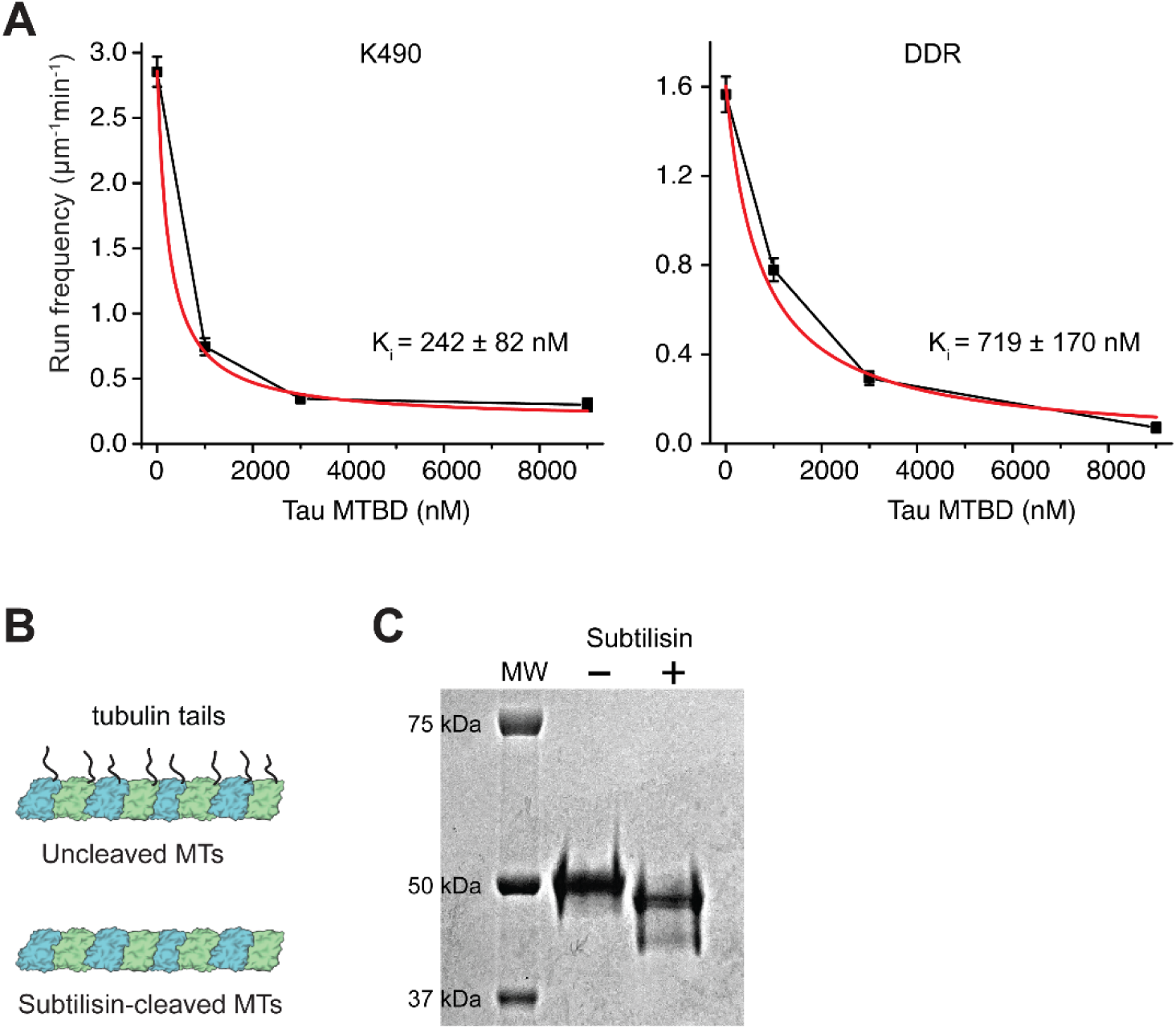
Tau MTBD inhibits both kinesin and dynein motility. **A.** Run frequency (mean ± SEM) of K490 and DDR was fitted to an inverse Langmuir isotherm (red curves) to calculate K_i_ (±SE). From left to right, n = 755, 583, 262, 77 runs for K560 and 445, 596, 167, 24 runs for DDR. **B.** Schematic of subtilisin cleavage of tubulin C-terminal tails. **C.** The denaturing gel shows a reduction in tubulin molecular weight (MW) upon subtilisin treatment.

**Fig. S4.**
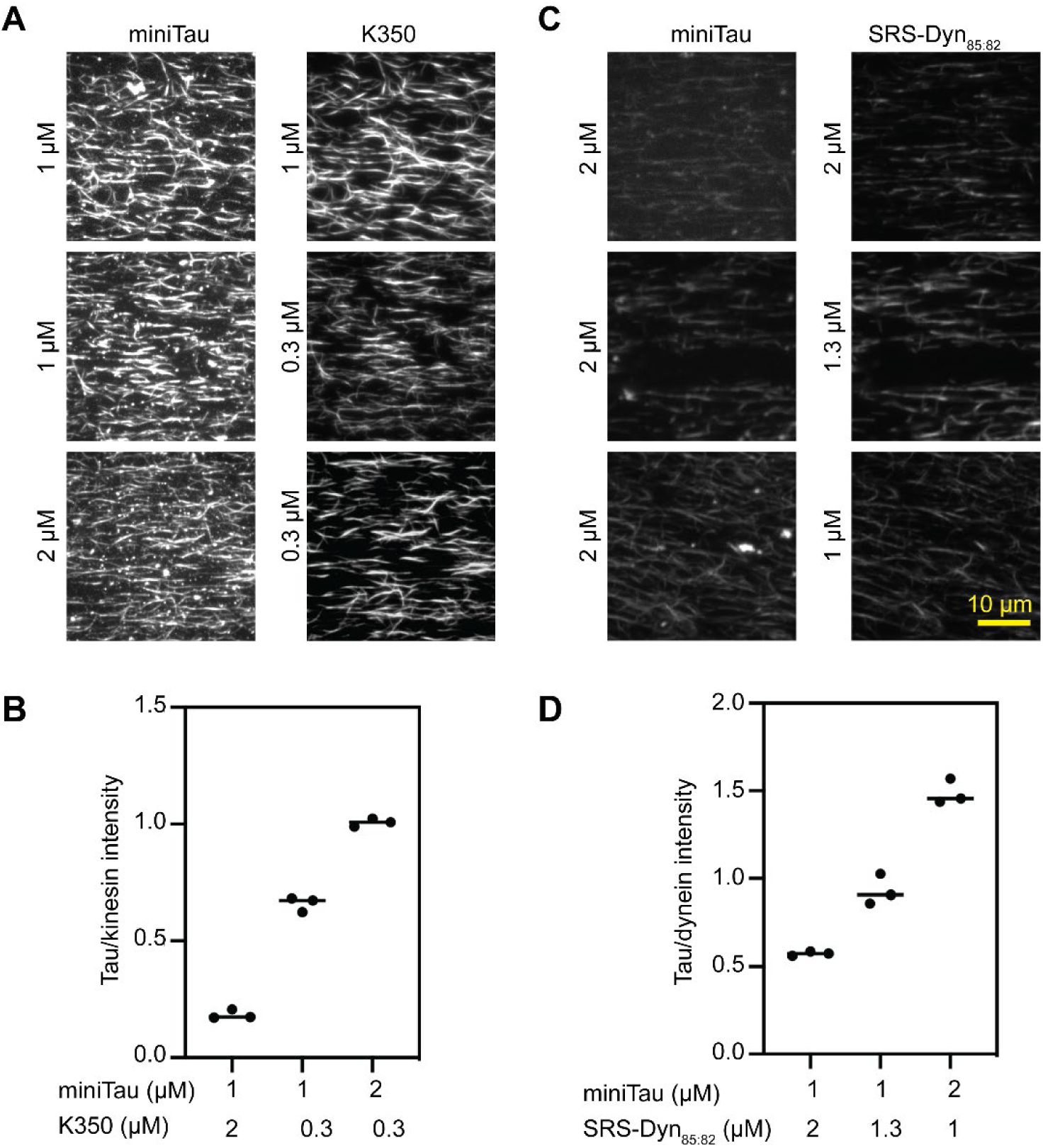
Optimization of the MT coverage of tau and motor proteins for cryo-EM studies. **A.** Example fluorescence images of MTs decorated with miniTau and K350^E236A^ at different concentrations. **B.** The ratio of the fluorescence intensities of miniTau to K350^E236A^ in three technical replicates. **C.** Example fluorescence images of MTs decorated with miniTau and SRS Dynein_85:82_ at different concentrations. **D.** The ratio of the fluorescence intensities of miniTau to SRS Dynein_85:82_ in three technical replicates. In B and D, the centerline represents the mean. The fluorescence intensity of each construct was normalized to its saturation condition (1 µM) on the MTs in the absence of other proteins before calculating the ratio of tau intensity to that of a motor protein.

**Fig. S5.**
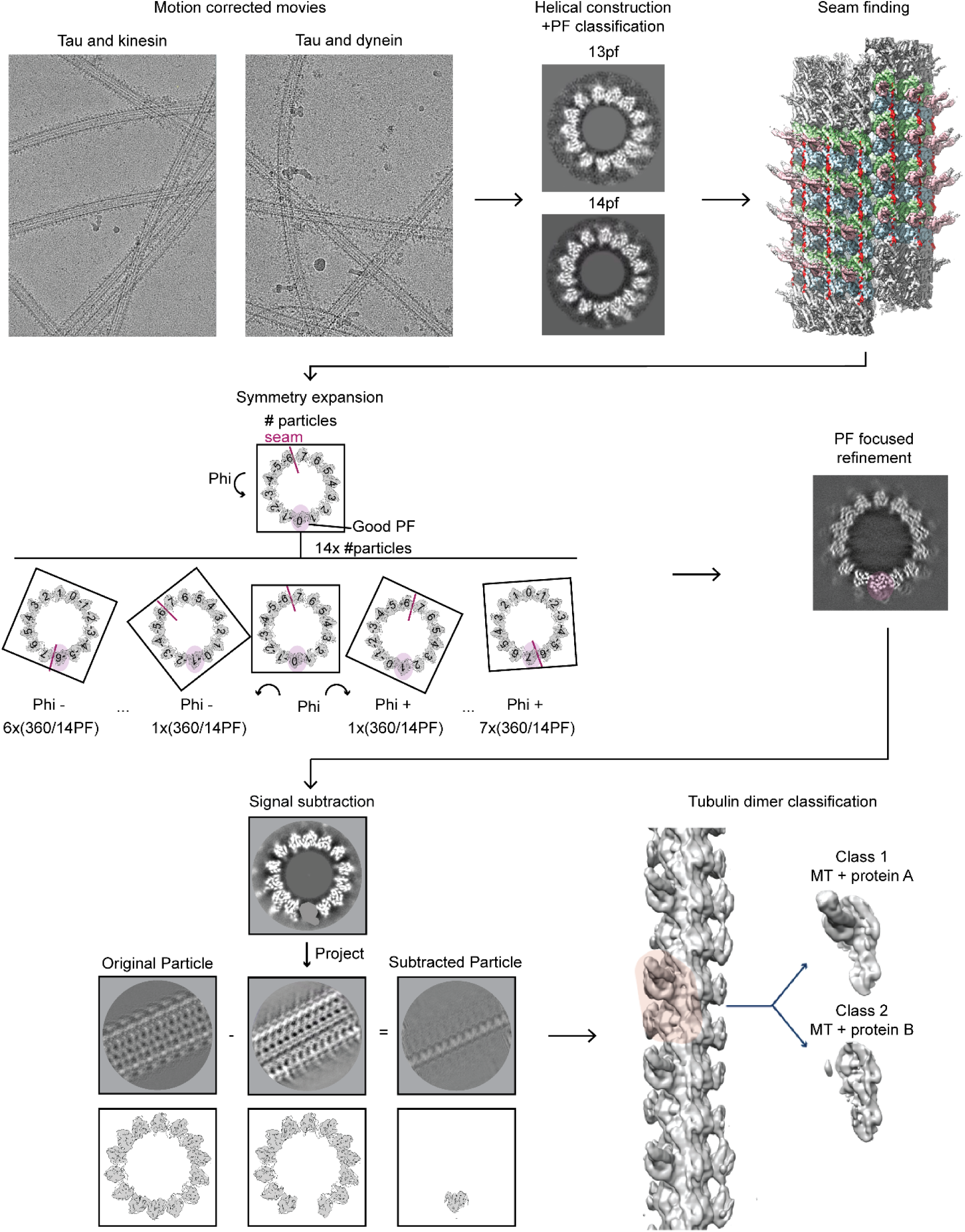
Cryo-EM data processing pipeline used to determine the co-structures of tau and a motor bound to MTs.

**Fig. S6.**
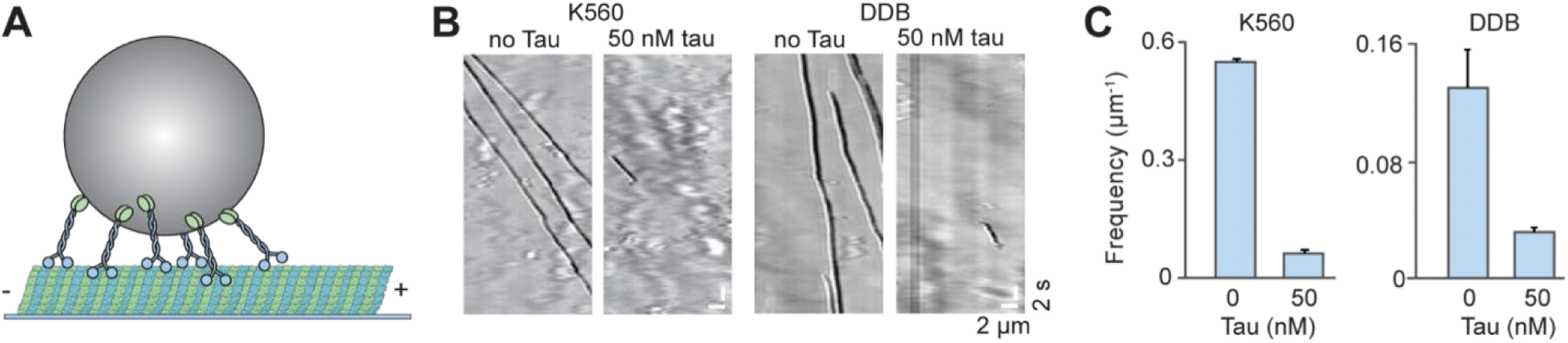
Tau inhibits beads driven by multiple kinesin motors. **A.** The schematic shows a bead driven by multiple kinesin motors. **B.** Beads driven by multiple K560s or DDBs are inhibited by 50 nM tau in the absence of added salt or methylcellulose. **C.** The run frequency for multi-motor-driven beads decreases with tau. From left to right, N = 210, 41, 109, and 28 runs. Error bars represent s.d.

**Fig. S7.**
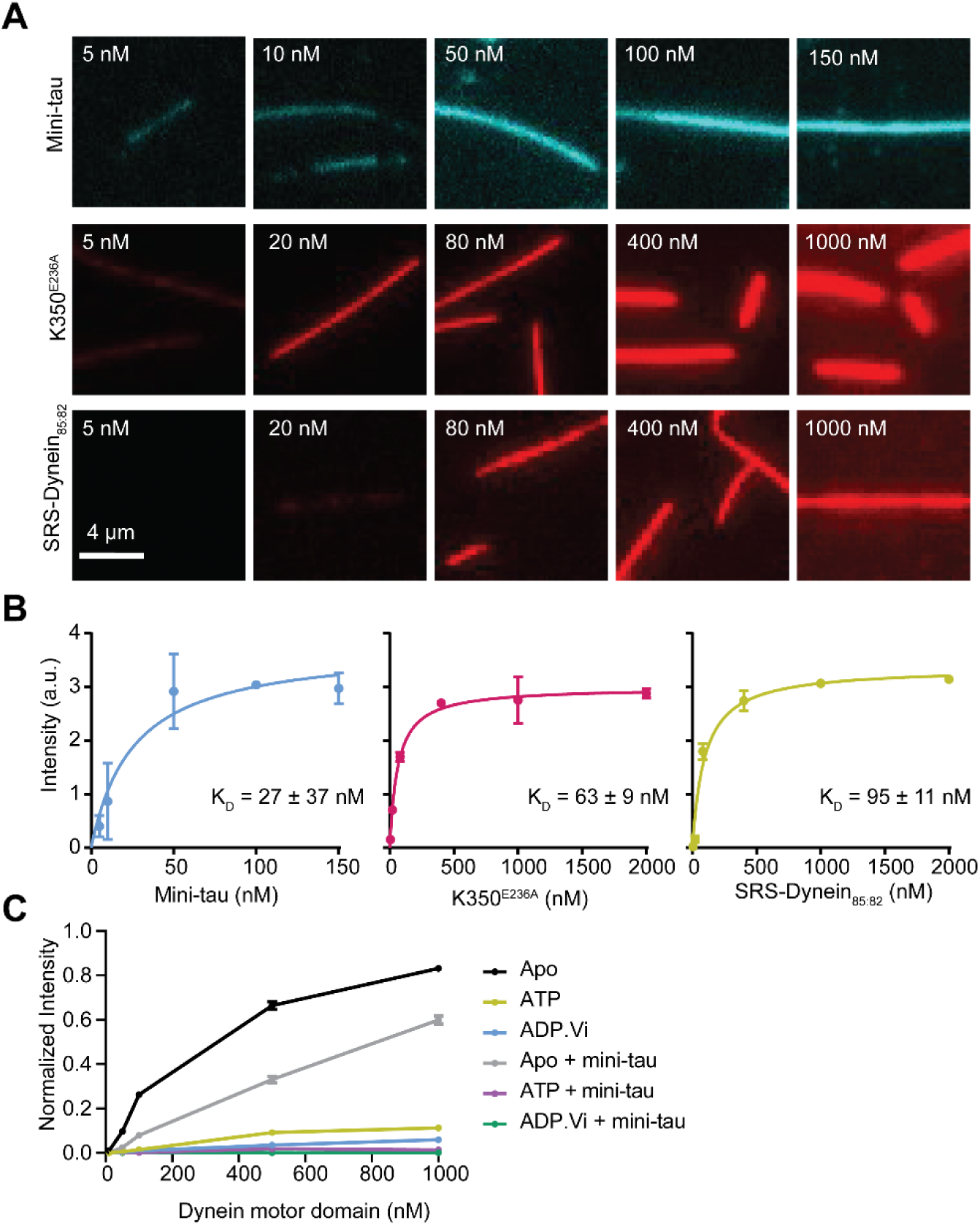
MT binding of dynein, kinesin, and mini-tau measured using fluorescence imaging. **A.** Representative images show MT decoration by K350^E236A^, SRS-Dynein_85:82,_ and mini-tau at different concentrations**. B.** The normalized intensities of K350^E236A^, SRS-Dynein_85:82_, and mini-tau binding to MT (mean ± S.D.). The assay was performed in BRB80 buffer without added salt or methylcellulose. From left to right, n = 191, 136, 204, 105, 227, 206 for K350^E236A^; and 126, 130, 167, 205, 227, 118 for SRS-Dynein_85:82_; and 170, 172, 158, 171, 104 for mini-tau. The dissociation constant (K_D_, ± S.D.) was calculated from a fit to the Hill Equation (solid curves). **C.** Normalized intensities of the dynein motor domain in the presence or absence of 100 nM mini-tau (mean ± s.e.m.) at different nucleotide states. From left to right, n = 134, 250, 172, 113, 202 in the apo state; 117, 127, 159. 107, 168 in ATP; 53, 78, 51, 65, 78 in ADP.Vi; 126, 171, 161, 90, 138 in the apo state with mini-tau; 110, 91, 121, 155, 182 in ATP with mini-tau; and 63, 75, 106, 71, 69 in ADP.Vi with mini-tau.

**Fig. S8.**
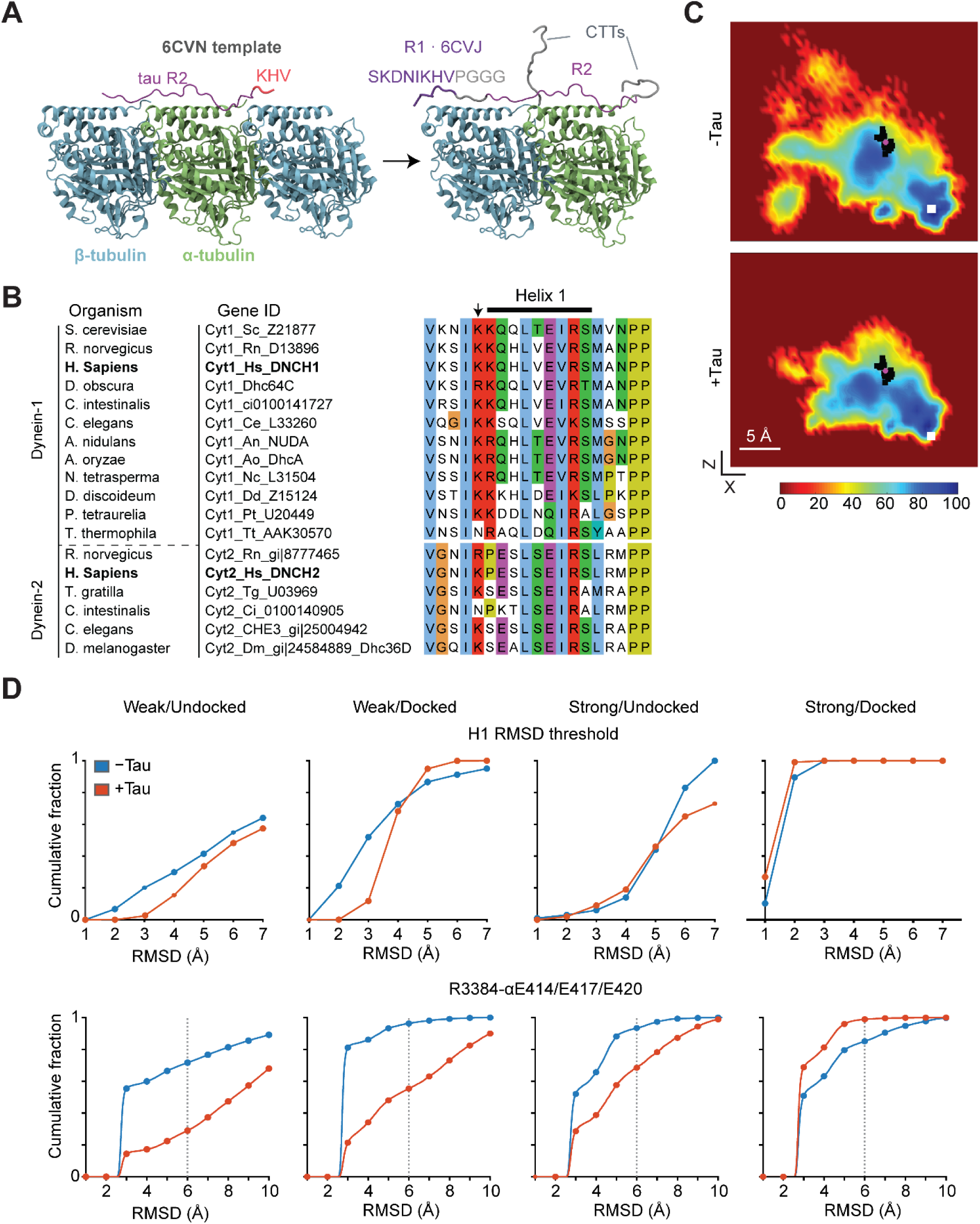
MD simulations of dynein MTBD in the presence and absence of the R2 sequence of tau. **A.** The structure of the R2 repeat of tau on βαβ tubulin (PDB: 6CVN) was used as a template for building the atomic model. For tau to span a tubulin heterodimer, β-tubulin located at the C-terminus of the tau structure and tau residues KHV were removed, and the tau R2 structure was extended by adding residues SKDNIKHVPGGG to its N-terminal end. Tubulin CTTs were added as intrinsically disordered peptides. **B.** Alignment of dynein-1 (Cyt1) and dynein-2 (Cyt2) heavy chain amino acid sequences from different organisms. The arrow points to the conserved lysine residue (K2996 in human dynein-2 and K3298 in human dynein-1) at the N-terminus of H1. **C.** The heat maps show H1 sampling pooled across the full trajectories of 25 simulations per condition (250 ns each; 6,250 ns total per condition). Black squares show the XZ position of the H1 center of mass for individual trajectories. The pink circle shows the average position of the black squares. White squares show the expected position of the H1 center of mass on tubulin at the strongly bound state of dynein. **D.** Cumulative fraction of the RMSD of H1 from its predicted docking site (top) and between dynein R3384 and E414, E417, and E 420 of α-tubulin.

## Methods

### Protein purification

Human FL tau from Sf9 cells, kinesin-1 heavy chain KIF5B (K560: amino acids 1-560 and K490: amino acids 1-490), phi mutant dynein, yeast dynein motor domain (Dyn_331kDa_)^43^, mouse BicD2N and mouse BicDR1 constructs have an N-terminal ZZ affinity tag, followed by a TEV protease cleavage site. The monomeric kinesin (K350: amino acids 1-350) and SRS-Dynein_85:82_ constructs have a 6XHis tag on the C-terminus. Human FL tau expressed in *E. coli*, and tau MTBD and Mini-tau expressed in insect cells have a Strep tag II on their N-terminus. The constructs used in this study are listed in Table S1.

FL tau, Mini-tau, tau-MTBD, K350^E236A^, and SRS-Dynein_85:82_ constructs were transformed into BL21 competent cells. Protein expression was induced with 0.5 mM IPTG, followed by 16 h incubation of the cell culture at 20 °C. The other constructs were transformed into DH10Bac competent cells, and the bacmids were purified and transfected into SF9 cells to make the virus. SF9 cells were passaged twice for 3 days at 27 °C to make the p2 virus. 10 mL of p2 was then added to 1 L of suspended SF9 cells and allowed to incubate for 3 days at 27 °C. Cells were pelleted at 4,000 g for 10 min in a JLA 8.1000 rotor.

To extract proteins, a 500 mL cell pellet for all constructs was re-suspended in 50 mL of either lysis buffer A for FL tau, mini-tau, tau-MTBD, K350^E236A^, and SRS-Dynein_85:82_ (50 mM Tris-HCl, pH 7.5, 1M potassium acetate, 10 mM MgCl2, 1mM EGTA, 10% glycerol, 1 mM PMSF, 1 mM DTT, and 1 tablet of protease inhibitor (Sigma), or lysis buffer B for K560, K490, phi mutant of dynein, dynein motor domain, and BicDR1 (25 mM HEPES pH 7.4, 1 M KAc, 1 mM DTT, 10% glycerol, 1 mM PMSF, and 1 tablet of protease inhibitor cocktail). BL21 cells were lysed with a tip sonicator for 2 min of active time. SF9 cells were lysed with a Wheaton glass dounce homogenizer. The lysate was clarified by spinning at 360,000 g for 45 min in a Ti70 rotor. The supernatant was incubated with 3 mL IgG beads, StrepXT beads, or Ni-NTA HisPure beads for constructs tagged with a Zz tag, Strep tag II, or 6XHis tag, respectively, for 2 h at 4 °C. Beads were then washed with 20 mL lysis buffer and 30 mL of either wash buffer A (50 mM Tris-HCl, pH 7.5, 10 mM MgCl2, 1mM EGTA, 10% glycerol, 1 mM DTT) or wash buffer B (25 mM HEPES pH 7.4, 150 mM KAc, 2 mM MgCl_2_, 1 mM EGTA, 1 mM PMSF, 1 mM DTT, and 1 tablet of protease inhibitor cocktail). Proteins with a Zz tag were eluted from IgG beads by incubating with 60 µL of 0.1 mg/mL TEV protease for 1 h at room temperature. Proteins with a Strep-tag II were eluted with 100 mM d-biotin. His-tagged proteins were eluted by adding 500 mM imidazole to the wash buffer. Six fractions of 1 mL were collected for each protein. Lysis and wash buffers contained 20 mM imidazole for proteins with a 6XHis tag and 1 mM ATP for motor proteins. For proteins labeled with a maleimide-reactive dye, TCEP was used instead of DTT in all buffers.

For fluorescent labeling, proteins were incubated with 4-fold excess dye derivatized with benzylguanine (BG, for SNAP labeling), coenzyme A (CoA, for ybbR labeling), maleimide (for K350^E236A^ and K490 CLM), or NHS (for SRS-Dynein_85:82_) at room temperature for 1 h. 5 µM SFP enzyme was added to catalyze ybbR labeling with CoA. Labeled proteins in the elution buffer were further purified to remove excess dye and other impurities. Dynein was eluted from the TSKgel G4000SWXL column (Tosoh), BicD2N, BicDR1, and K490 were eluted from the Superose 6 10/300 GL column (Cytiva), and sf9 FL tau and K560 were eluted from the Superdex 200 Increase 10/300 GL column (Cytiva). FL tau expressed in E.coli was eluted from a Superose 6 10/300 GL column (Cytiva) as previously reported^17^. Mini-tau was applied onto a 6 mL Resource S column (Cytiva) and eluted with a gradient of lysis buffer. Desired fractions were collected and concentrated with an Amicon Ultra centrifugal filter.

Dynactin was purified from pig brains using SP Sepharose Fast Flow, MonoQ ion exchange columns (Cytiva), and a TSKgel G4000SWXL size exclusion column (Tosoh)^52^.

### MT polymerization

Tubulin was purified from pig brains and labeled with biotin or fluorophores. Biotinylated MTs were polymerized by mixing 4 μL of 3.5 mg/mL biotinylated tubulin and 5 μL of 25 mg/mL unlabeled tubulin in BRB80 buffer (80 mM PIPES pH 6.8, 1 mM MgCl_2_, 1 mM EGTA) with 100 μM taxol at 37 °C. LD555-labeled MTs were polymerized under the same conditions with 1 μL of 2 mg/mL LD555-labeled tubulin.

### Preparation of coverslips and flow chambers

Glass coverslips were functionalized with PEG/PEG-biotin to reduce the nonspecific binding of proteins. Plain glass coverslips were cleaned sequentially with water, acetone, and water by sonication for 10 min, and then sonicated in 1 M KOH with a bath sonicator for 40 min. The coverslips were then incubated in a mixture of 3-Aminopropyltriethoxysilane, acetic acid, and methanol with a ratio of 1:5:100 for 10 min, with 1 min sonication between two successive incubations. 30 µl of 25% biotin-PEG-succinimidyl valerate in a 0.1 M NaHCO_3_ buffer (pH 8.5) was sandwiched between two pieces of air-dried coverslips after being washed with methanol and allowed to incubate at 4 °C overnight. The sandwiches were separated the next day and washed with water. The biotinylated coverslips were air-dried and kept at −20 °C under vacuum conditions for long-term storage. The coverslips were later assembled into 10 µL flow chambers with laser-cut double-sided tape.

### Fluorescence Microscopy

Fluorescence microscopy was performed using a Ti-E Eclipse inverted microscope with a 100x 1.49 NA Apo TIRF oil immersion objective and a TIRF Motorized Illuminator unit (Nikon). Samples were excited using 488, 561, and 633 nm lasers (Coherent) fiber-coupled through a single-mode fiber via a wave-division demultiplexer (Oz Optics). Emission signals were filtered using 512/25, 580/40, and 697/75 nm bandpass filters (Chroma) mounted in a Lambda 10-B optical filter changer (Sutter). Images were captured with an EMCCD camera (Andor, iXon+). The microscope was controlled using Andor Solis and MicroManager for stepping and single-molecule motility experiments, respectively. The effective pixel size was set to 86.7 nm and 160 nm after magnification for stepping and motility experiments, respectively.

### Single-molecule motility assays

Biotinylated MTs were diluted in MB (30 mM HEPES pH 7.4, 5 mM MgSO_4_, 1 mM EGTA, 1 mg/mL casein, 0.5% pluronic, 1 mM TCEP, and 10 µM taxol) to a final concentration of 20 nM polymerized tubulin heterodimer. 10 µL streptavidin at 5 mg/mL was added to the chamber, incubated for 2 min, and washed twice with 20 µL MB buffer. The chamber was then incubated with 10 µL diluted MTs for 2 min and washed with 20 µL MB buffer to remove unbound MTs. Proteins were diluted and mixed into desired concentrations in MB buffer. For DDR assembly, 20 nM dynein, 150 nM dynactin, and 50 nM BicDR1 were pre-mixed on ice for 30 min, then diluted 7-fold using MB buffer. The final protein mixture was diluted in the imaging buffer (MB buffer supplemented with 150 mM KAc, 0.1% methylcellulose, 2 mM ATP, 2 mg/mL glucose oxidase, 0.5 mg/mL catalase, 0.8% glucose, motor proteins, and tau). The sample was sealed and imaged immediately. Reported protein concentrations represent the final concentrations in the chamber. Each movie was recorded for at least 150 frames per imaging area. MTs were visually identified using the z-stack feature of ImageJ to generate kymographs. Kymographs were analyzed using a custom-written MATLAB code for run frequency, run length, and velocity calculations.

### MT binding assay and data analysis

PEG/PEG-biotin surfaces were incubated with 10 µL of 5 mg/mL streptavidin for 2 min. The chamber was washed twice with 10 µL BRB80CPT buffer (80 mM PIPES pH 6.8, 10 mM MgCl_2_, 5 mM EGTA, 1 mM DTT, 1 mg/mL casein, 0.5% pluronic, and 1 µM taxol). Biotinylated MTs were diluted in BRB80CPT to a final concentration of 20 nM polymerized tubulin heterodimer and introduced to the chamber for 3 min. The chamber was washed with 10 µL BRB80CPT twice to remove unbound MTs. Proteins were diluted to the desired concentration in 10 µL of imaging buffer (BRB80CPT buffer supplemented with 2 mg/mL glucose oxidase, 0.5 mg/mL catalase, 0.8% glucose) and flowed into the chamber. The chamber was imaged immediately. Average signals from MTs were identified and calculated manually using ImageJ. For assays with the dynein motor domain, the catalytically dead motor population was calculated by washing the chamber once with 20 µL BRB80CPT and twice with 20 µL of BRB80CPT supplemented with 2 mM ATP and 2 mM sodium vanadate. The remaining fluorescence signal on the MT was subtracted from the raw signals in other nucleotide conditions. The run frequency (RF) of K490, K560, and DDR in different tau concentrations was fitted to a Langmuir isotherm in its inhibitory form 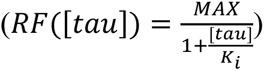 where *MAX* and *MIN* are the maximum and minimum run frequencies, and *K_i_* is the inhibition constant.

### Stepping assays

A cysteine-light mutant of K490 E215C was labeled at the motor domain with biotin-maleimide. Biotinylated-K490 was diluted to the desired concentration in BRB80CPT, and 9 µL of K490 was mixed with 1 µL of 1 µM streptavidin-Qdot 655 (Invitrogen) for 3 min. MTs were diluted and immobilized on the biotinylated coverslips as described above. 1 µL of 1 mM d-biotin was added to the K490-QDots mixture for 5 min to biotinylate streptavidin. 1 µL of tau at a desired concentration was then introduced to the same mixture. The chamber was washed with 10 µL BRB80CPT buffer twice and then with 20 µL of 1 mM d-biotin. The protein mixture was flown into the chamber and incubated for 5 min. 10 µL of stepping buffer (BRB80CPT buffer supplemented with 2 mg/mL glucose oxidase, 0.5 mg/mL catalase, 0.8% glucose, and 500 nM ATP) was then introduced into the chamber. Movies were acquired at 76 ms per frame for 1400 frames. The diffraction-limited spots of individual motors were localized and tracked by a modified version of FIESTA^53^. The trajectories of single motors were fitted to steps in MATLAB using a Schwartz information criterion algorithm^54^. Fitted traces were manually inspected and corrected for missed steps.

### Cryo-EM sample preparation

Porcine brain tubulin (Cytoskeleton) was reconstituted in BRB80 buffer (80 mM PIPES, pH 6.9, 1 mM EGTA, 1 mM MgCl_2_) supplemented with 10% glycerol and 1 mM GTP, clarified by ultracentrifugation, and snap-frozen for storage at −80 °C. MTs were polymerized in the presence of peloruside A (PelA)^12^. Tubulin aliquots containing 1 mM GTP were incubated at 37 °C with two sequential additions of 200 µM PelA for 20 min, then pelleted at 37 °C at 15,000 rpm for 20 min. The MT pellet was resuspended in EM buffer (80 mM PIPES, pH 6.8, 1 mM EGTA, 1 mM MgCl_2_, 1 mM DTT, 0.05% Nonidet P-40) containing 250 µM PelA. MT concentration was determined on a NanoDrop spectrophotometer by measuring OD_280 nm_ of a depolymerized aliquot incubated with 0.1 M CaCl_2_ for 10 min on ice. MTs were then diluted to the desired concentration for grid preparation in EM buffer containing 100 µM PelA.

To determine whether and how tau and kinesin or dynein can co-occupy the same tubulin dimer, cryo-EM grids of PelA-MTs decorated with mini-tau and either K350^E236A^ or SRS-Dynein_85:82_ were prepared by diluting these constructs in EM buffer prior to combining with MTs. To account for their different MT affinities and to increase the likelihood of tau and a motor protein occupying the same tubulin, TIRF imaging was used to test different concentrations of tau and motor protein to achieve similar, near-saturating occupancy of both proteins on the MT lattice. Tau was applied to the MTs first, followed by a mixture of tau and the respective motor protein. Protein concentrations and molar ratios were subsequently optimized by screening a range of conditions by cryo-EM to account for differences in sample preparation for cryo-EM. Conditions were selected in which both proteins were readily detected at comparable densities in initial reconstructions of MT segments.

For the tau–dynein sample, 2.5 µl of 2.5 µM PelA-MTs was applied for 30 s to a glow-discharged Au300 R1.2/1.3 holey-carbon grid and manually blotted. This was followed by sequential application of 2.5 µl of 1 µM mini-tau for 30 s and manual blotting. Finally, 2.5 µl of a freshly prepared mixture of 1 µM mini-tau and 3 µM SRS-Dynein_85:82_ was applied and incubated for at least 30 s while the grid was mounted on a Vitrobot Mark IV (ThermoFisher). The sample was then blotted for 7 s using a blot force of 6 and plunged into liquid ethane. The Vitrobot was operated at 20 °C and 80% humidity.

Tau–kinesin grids were prepared using a similar protocol, with optimized concentrations of 2 µM mini-tau and 1 µM K350^E236A^. Specifically, 2.5 µl of 2.5 µM PelA-MTs was incubated on a glow-discharged grid for 30 s and blotted. Then, 2.5 µl of 2 µM mini-tau was added, incubated for 30 s, and manually blotted. Finally, 2.5 µl of a freshly prepared mixture of 2 µM mini-tau and 1 µM K350^E236A^ was applied, and the grid was blotted for 5 s in the Vitrobot, using a blot force of 5.

### Cryo-EM data collection

Cryo-EM data were collected on a Talos Arctica microscope (ThermoFisher) operated at an accelerating voltage of 200 kV. Images were recorded on a K3 direct electron detector (Gatan) operated in super-resolution counting mode at a nominal magnification of 36,000×, corresponding to a calibrated physical pixel size of 1.14 Å (0.57 Å in super-resolution mode). Movies were collected at a dose rate of 5–8 electrons per pixel per second, for a total dose of ∼50 electrons Å^−2^ on the specimen. Data were collected semi-automatically using SerialEM. Cryo-EM dataset and model-building statistics are shown in Table S2.

### Cryo-EM data processing

Cryo-EM data of tau- and motor protein-bound MTs were processed as previously described^38^. Movie stacks were motion-corrected and dose-weighted using MotionCor2, and the contrast transfer function (CTF) of each micrograph was estimated using Gctf^55^. Micrographs with a poor CTF fit (estimated resolution lower than 6 Å) or a low figure of merit (below 0) were discarded. MTs were manually picked as helical filaments in RELION 3.0. Overlapping MT segments were initially extracted as binned particles (bin 2) with a box size of 256 binned pixels and a step size of 82 Å (the length of a tubulin dimer) along the filament axis. The protofilament (PF) number of each MT was determined by reference-based 3D classification against 13- and 14-PF references low-pass filtered to 20 Å. After classification, the PF class distribution of each helical filament was evaluated, and if more than 80% of the particles belonged to the same class, all particles from that segment were assigned to this class. Filaments with an inconclusive PF distribution were discarded. Particles from the dominant 14-PF class were selected for further processing.

Global alignment parameters of the selected particles were refined by 3D helical refinement in RELION 3.0. Helical rise and twist were then determined using the *relion_helix_toolbox* function and imposed as pseudo-helical symmetry constraints in the following steps. Particles were then extracted and unbinned with a box size of 512 pixels for subsequent processing.

To determine the position of the MT seam for each particle, unbinned particles were first imported into FREALIGN v9^56^ and refined, considering the pseudo-helical symmetry of a 14-PF microtubule and applying the previously determined helical parameters. The seam position of each particle was then determined as previously described^37^. Once the seam position and the unique phi angle around the MT axis had been determined for each particle, particles were further refined in FREALIGN v9.

To determine whether tau and the respective motor proteins occupy the same tubulin dimer or exclude each other from binding to the same tubulin dimer, we performed symmetry expansion, signal subtraction, and focused classification, as previously described^38^. Particles were symmetry expanded around a selected PF (the “good” PF) of each MT segment, followed by focused 3D refinement of the same PF in either RELION 3.0 or cisTEM^57^ using a soft mask around the PF. Signal subtraction was then performed in RELION 3.0 to isolate the density of the good PF and to facilitate classification of the small protein domains bound to the MT.

Using the signal-subtracted micrographs and the refined positions and orientations, focused 3D classification was performed without alignment in RELION 3.0, applying a soft mask around the MT-binding sites of the motor proteins. For kinesin, focused classification was performed with a soft mask around the kinesin motor domain, requesting two classes with a T value of 40^38^. Classifications with more classes yielded additional, nearly empty classes, indicating that most particles fall into these two classes. For dynein, because of the smaller footprint of the dynein MTBD, the mask was extended to include the full repeat of the tau MTBD. Classification of dynein and tau was performed in RELION 3.0, requesting two or four classes with a T value of 40.

### Rigid-body fitting of atomic models into cryo-EM maps

For interpretation and visualization of the maps before and after focused classification, existing structural models of the proteins were fitted as rigid bodies into the cryoEM maps using the fit-in-map function in UCSF Chimera. The respective models were first fitted into the single protofilament-focused refined maps (Fig. 3B, 3.5 Å resolution) obtained before tubulin dimer classification. The tau density was fitted with the R2 tau model from PDB ID: 6CVN^12^, which marks the tau footprint without assigning a specific repeat. A PelA-stabilized tubulin dimer (PDB ID: 5SYC)^48^ was then fitted into the tubulin density, which also accommodated the PelA density, and replaced the 6CVN tubulin while the tau model was retained. The fitted PelA-stabilized tubulin dimer and the tubulin dimer in the R2-tau bound model did not display substantial (root-mean-square deviation (RMSD) of Cα atoms was less than 1 Å) shifts relative to each other.

MT binding domains of kinesin and dynein were fitted into their respective densities. For dynein, the high-affinity MTBD–tubulin complex (PDB ID: 6RZB)^45^ was fitted, and the MTBD was then refitted as a separate rigid body, revealing a small visual change (Cα RMSD 2.8 Å) of the dynein position relative to the tubulin dimer. The obtained fit confirmed no steric clashes between the tubulin-bound tau model and the tubulin-bound dynein model. For kinesin, we used PDB ID: 3J8X, corresponding to the nucleotide-free K349 motor domain on MTs^47^. The motor domain was fitted together with tubulin and then refitted alone, showing no substantial shift (Cα RMSD = 0.5 Å). As expected, this fit revealed a steric clash between the tau model and the kinesin model. After generating the combined models from the maps, those models were compared to the class averages obtained from the tubulin dimer classification (Fig. 3C,D).

### Atomic model building for MD Simulations

Dynein motor domain conformations were sampled from the MD simulations performed on the ADP.Pi-bound structure of human dynein-2 in the absence of MTs^49^. The MTBD and parts of stalk coiled-coils 1 and 2 (CC1 and CC2, respectively) of human dynein-2 (amino acids 2962 – 3126) were used to investigate dynein binding to a tau-free or tau-bound tubulin heterodimer. The high-affinity dynein MTBD structure, along with its adjacent portions of CC1 and CC2, was taken from the mouse cytoplasmic dynein-1 MTBD cryo-EM structure resolved in complex with α/β-tubulin (PDB ID: 6RZB)^45^. Two alanine substitutions (A3323 and A3387) introduced in the cryo-EM construct were reverted to their native cysteine residues with the Mutator plugin in VMD. Because tubulin CTTs could not be resolved by cryo-EM, they were absent from the starting cryo-EM model. Residue numbers in Figures 6 and S8 are given for mouse dynein-1.

The cryo-EM structure of the R2 repeat of tau on βαβ tubulin (PDB: 6CVN)^12^ was used as a template for simulations. For a tau pseudorepeat to span a single αβ tubulin heterodimer, tau residues K298-V300 were removed, and residues S262-G273 were added to its N-terminal end based on the structure of the R1 repeat of tau (PDB ID: 6CVJ)^12^. The missing region comprising residues P270-G273 was constructed using the Molefacture plugin of VMD. The missing CTTs of tubulin were first built as intrinsically disordered peptides using the Molefacture plugin and structurally optimized through 200 ns-long MD simulations. The resulting CTT conformations were added to the α/β-tubulin model by orienting them towards the solvent while avoiding steric clashes.

To dock the dynein conformations to the tau-tubulin complexes, the high-affinity dynein-1 MTBD–tubulin complex (PDB ID: 6RZB) was superimposed onto the tau–tubulin complex (PDB ID: 6CVN) using the C_α_ atoms of the secondary structure elements of α- and β-tubulin. Then, the low-affinity dynein-2 conformations were superimposed on the dynein-1 MTBD using dynein MTBD helices 1, 2, 3, and 6. To model dynein binding to tubulin, high- and low-affinity dynein starting structures were generated by shifting them 7 Å away from the dynein–tubulin interface along the tubulin surface normal. The tau-free systems were generated by removing tau from each system.

### All-atom MD simulations

Each system was solvated in a water box using the TIP3P water model with padding of at least 15 Å of water in each direction. The solution was ionized to 150 mM NaCl. Each system was composed of ∼200,000 atoms. MD simulations were performed in NAMD 3 using the CHARMM36m all-atom additive protein force field, as described previously^58^. For tau systems, dynein and tubulin were kept fixed for 10,000 steps of minimization, followed by 1 ns equilibration. Next, constraints on dynein and tubulin were released for an additional 10,000 steps. Subsequently, 2 ns of equilibration were performed, during which harmonic potentials having a spring constant of 1 kcal mol^−1^Å^−2^ were applied to the C_α_ atoms. Before initiating the MD simulations, all constraints were removed, and the system was equilibrated for an additional 4 ns. In tau-bound simulations, harmonic constraints were applied to C_α_ atoms of tau and the tubulin residues (Table S3A). In total, 68 runs of MD simulations were carried out for dynein-1 and dynein-2 systems, covering tau-bound, tau-free, and 7 Å distant (encounter-state) configurations (Table S3B). Each simulation was performed for 250 ns, yielding an aggregate simulation time of 17 μs.

Principal axes (PA) of MTs were obtained via the orient tool in VMD. PA1 corresponds to the long axis of a protofilament. PA2 (short axis) is the vector orthogonal to PA1 that passes through the center of mass (CoM) of the Q2982, K2966, Y3099, and E3117 Cα atoms of human dynein-2. The x and z coordinates of the H1 CoM were calculated and projected onto the plane constructed by PA1 and PA2. Subsequently, heat maps were constructed from the density distribution of these projections in MATLAB. The stalk coiled coil vector, defined as the vector pointing from the Q2982/Y3099 CoM to the K2966/E3117 CoM, was projected onto the PA1-PA2 plane. The angle between the projected vector and PA2 was defined as the stalk angle. The H1 vector (S3005 Cα → E2998 Cα of human dynein-2) was analyzed similarly to obtain its angular orientation. Salt bridges were identified using a 6 Å cutoff between basic nitrogens and acidic oxygens,^59^ while hydrogen bonds were defined by donor–acceptor distances ≤ 3.5 Å and donor–H–acceptor angles ≤ 30° ^60^. Only conformations with an H1 RMSD equal or below 0.5 nm relative to the docked reference structure were included in analyses.

### Statistical analysis

*P* values were determined using statistical tests in Prism. Two-sided two-sample t-tests were performed to compare continuous datasets, assuming that data are approximately normally distributed. Each result was based on a minimum of two independent experiments. The number of replicates and data points (n) and the specific statistical methods used are provided in the figure legends. Representative data from independently repeated experiments are shown.

## Data Availability

A reporting summary for this article is available as a Supplementary Information file. The main data supporting the findings of this study are available within the article and its Extended Data Figures. Protocols supporting the findings of this study are described in the Methods section. The constructs that express WT and mutant versions of tau will be deposited at AddGene. Raw microscopy data will be made available by the corresponding authors upon request.

## Code availability

The custom code used to analyze experimental data is uploaded to Zenodo (doi: 10.5281/zenodo.18041601). Scripts for cryo-EM image processing are available from E.N. upon request.

## Movie captions

**Movie S1. K560 and DDR motility on MTs in the presence of different tau concentrations** K560 motors were labeled with LD655-BG on a C-terminal SNAP-tag. The DDR complex was labeled with LD655-BG on a C-terminal SNAP-tag of BicDR-1. Tau is added at the given concentration along with the motor solution and is not washed out of the chamber. Motility buffer includes 150 mM KAc, 0.1% methylcellulose, glucose oxidase, catalase, dextrose, and 1 mM ATP. K560 and DDR motility were recorded at 5 Hz under TIRF illumination. MT decoration on the PEG-biotin coverslip was recorded in a separate fluorescent channel (not shown).

**Movie S2. Binding of dynein to the MT in the presence and absence of the R2 repeat of tau on tubulin** Time-lapse movies of MD trajectories show that a salt bridge between K2996 of dynein (corresponding to K3298 in human dynein-1) and E431 of β-tubulin facilitates docking of the H1 helix to its binding site. The formation of this salt bridge was interrupted by interactions between the backbone of the R2 repeat of tau and the dynein MTBD, initiated from the weak-affinity conformation, and released 7 Å away from the tubulin binding site.

**Movie S3. Tau affects the orientation of dynein on the MT** Time-lapse movies of MD trajectories show that, in the presence of tau, dynein MTBD initiated from the weak-affinity conformation and released 7 Å away from the tubulin binding site (yellow) is tilted away from tau by ∼10° in comparison to its stable docking orientation (grey) in the absence of tau.

